# Rewiring oncogenic signaling to RNA vector replication for the treatment of metastatic cancer

**DOI:** 10.1101/2024.12.30.630713

**Authors:** Xinzhi Zou, Cynthia Zhao, Kevin T. Beier, Chil-Yong Kang, Michael Z. Lin

## Abstract

Despite recent advances, improvements to long-term survival in metastatic carcinomas, such as pancreatic or ovarian cancer, remain limited. Current therapies suppress growth-promoting biochemical signals, ablate cells expressing tumor-associated antigens, or promote adaptive immunity to tumor neoantigens. However, these approaches are limited by toxicity to normal cells using the same signaling pathways or expressing the same antigens, or by the low frequency of neoantigens in most carcinomas. Here, we report a fundamentally different strategy for designing safer and more effective anti-cancer therapies through the sensing of cancer-driving biochemical signals and their rewiring to virotherapeutic activation. Specifically, we rationally engineer a RNA vector to self-replicate and cause cytotoxicity in cancer cells exhibiting hyperactive HER2 (ErbB2), but not in normal cells with normal HER2 signaling. Compared to a widely tested virotherapeutic from the same vector family, our hyperactive ErbB2-restricted RNA vector (HERV) exhibits lower toxicity and greater activity against metastatic HER2-positive ovarian cancer in mice, extending survival independently of tumor antigenicity. Most importantly, HERV synergizes with standard-of-care chemotherapy against ovarian cancer metastases in vivo, with 43% of combination-treated subjects surviving for months beyond subjects treated with either therapy alone. Taken together, these results introduce rewiring of cancer-driving signaling pathways to virotherapeutic activation as a strategy for more specific and effective cancer treatment.

## Introduction

Cancer is the leading cause of death in East Asia, Europe, and North America, and the second leading cause globally^1^. More than 90% of cancer deaths occur from metastatic or disseminated disease^2^, with epithelial tumors (carcinomas) such as lung, colon, liver, stomach, and breast cancer responsible for the largest loss of life years^3^. Standard of care in newly diagnosed or recurrent metastatic carcinomas generally consists of cytoreductive surgery followed by chemotherapy and/or therapies targeting specific cancer-associated proteins. However, long-term survival rates for metastatic carcinoma remain low; for example 5-year survival in metastatic lung and breast cancers, among the most actively studied metastatic cancers, remains below 30%^4,5^. Thus there is a large unmet need for new therapeutic entities that can further improve patient outcomes over current treatment regimens.

Recent advances in treating metastatic carcinomas have come from targeting tumor-driving signals, but benefits are still limited in degree and generalizability^6^. Constitutive phosphorylation of ErbB-family receptor tyrosine kinases (RTKs), which include ErbB1 (also named EGFR or HER1) and ErbB2 (also named HER2 or Neu) occurs in up to 28% of carcinoma cases^7,8^, and is associated with poor prognosis in breast and ovarian cancers^9,10^. Constitutive phosphorylation occurs either as a result of dimerization-activating mutations or gene amplification resulting in overexpression. Monoclonal antibodies (mAbs) targeting HER2 extend survival in metastatic HER2-overexpressing breast cancers, with the addition of the mAbs pertuzumab and trastuzumab to docetaxel chemotherapy now established as first-line therapy, and the antibody-drug conjugate (ADC) T-DXd established as monotherapy in second-line therapy^11–13^. T-DXd monotherapy also induces tumor responses in high-HER2 gastric and ovarian cancers^14,15^. However, median survival after treatment remains less than 5 years for breast cancer^11^ and 2 years for gastric and ovarian cancers, despite the ability to continue treatment with ADCs indefinitely^14,15^. Thus, despite recent advances, there remains substantial need to improve long-term survival from metastatic carcinomas. Small-molecule drugs that inhibit the tyrosine kinase activities of EGFR or HER2 are also efficacious in particular settings, but limited in benefit. EGFR tyrosine kinase inhibitors (TKIs) extend survival in metastatic EGFR-mutated lung cancer better than chemotherapy, but median survival remains less than 4 years^16^. HER2 kinase inhibitors improved survival in metastatic breast cancer when added to first-generation anti-HER2 antibodies, but median survival was still less than 2 years^17^, and their use has been superseded by recent ADCs. In addition, TKIs have not shown clear efficacy in HER2-positive ovarian cancer^18^ or gastric cancer^19,20^.

A major limitation of the above targeted therapies is the presence of dose-limiting on-target off-tumor toxicity. For TKIs and antibodies targeting EGFR or HER2, all receptor molecules are targeted whether they are in normal or cancer cells. As normal cells also express EGFR or HER2 and rely on their signaling for survival, the efficacious doses of these targeted agents overlaps with their toxic doses^21–23^. Thus TKIs cannot fully inhibit EGFR or HER2 signaling, and antibodies cannot fully ablate EGFR- or HER2-expressing cells, without causing toxicity to healthy tissues. Indeed CAR-T cells directed against HER2, which are able to expand *in vivo*, have shown unacceptably high levels of toxicity^24^.

We sought to design a new class of direct tumor-killing agents that can be highly cancer-specific and that use a different mechanism of action from existing therapies, allowing them to be combined to improve survival. We reasoned that if we could activate a therapy specifically within ErbB-hyperactive cells and not ErbB-normal cells, then it should be much less toxic than therapies that inhibit all ErbB signaling or ablate all ErbB-expressing cells. Previously, a system termed Rewiring Aberrant Signaling to Effector Release (RASER) was developed to rewire ErbB hyperactivity to the release of a programmable effector protein^25^. The ErbB-RASER system consists of two protein components: First, a releaser protein composed of a phosphoErbB-binding domain fused to a sequence-specific viral protease, and second, a substrate protein composed of a membrane tether, another phosphoErbB-binding domain, a protease-cleavable sequence, and a therapeutic effector domain **(Fig. 1a)**. The two proteins bind to activated ErbB receptors simultaneously, whereupon proximity-enhanced proteolysis releases the effector domain. Because release continues as long as active receptors are present, cancer cells with constitutively active EGFR or HER2 accumulate released effector to high levels^25^. In contrast, EGFR or HER2 activation by growth factors in normal cells is time-limited by receptor downregulation^26^, resulting in distinctly lower amounts of released effector^25^. By changing the effector domain, ErbB-RASER can be programmed to induce a variety of outputs including apoptosis and CRISPR/Cas9-mediated activation of endogenous cytokine genes^25^.

**Fig. 1.**
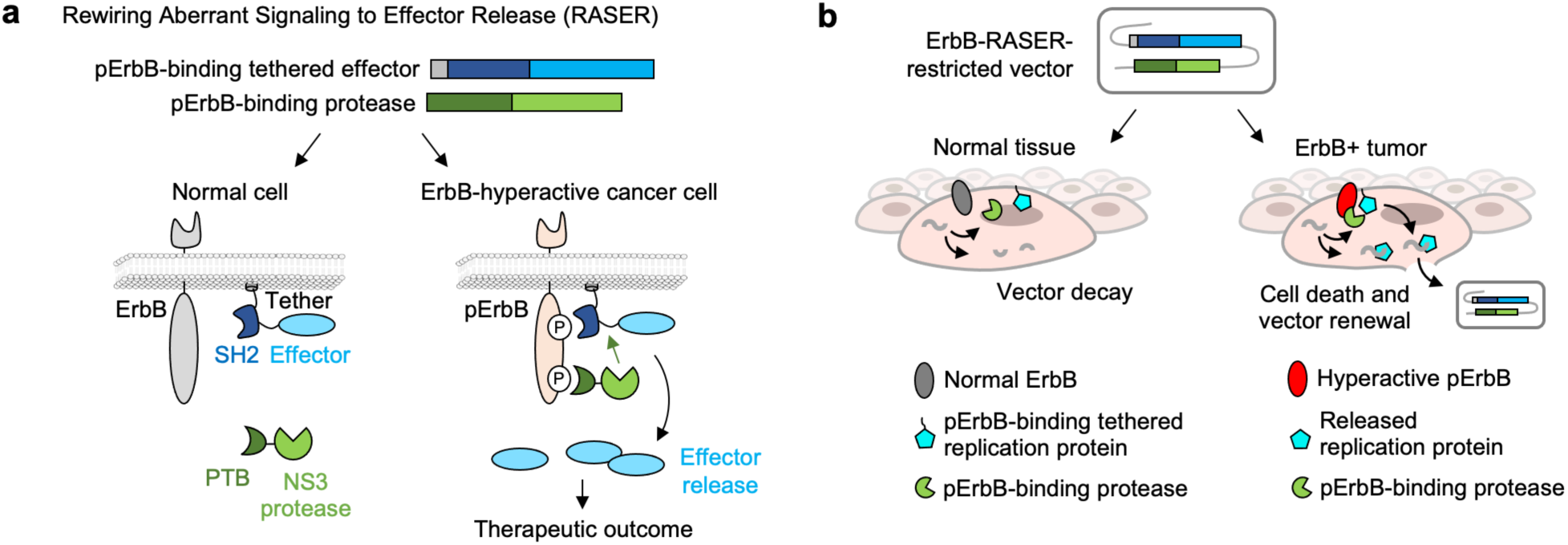
Concept for a ErbB-selective virotherapy using RASER. **a,** The ErbB RASER system consists of a pErbB-binding tethered payload and a pErbB-binding releaser. Effector is tethered on plasma membrane and will be released in an ErbB-activity dependent manner. **b,** Concept of using RASER to enhance viral vector safety by restricting viral replication in ErbB+ cancer cells. In normal cells, RASER vector could enter the cells but not replicate themselves, whereas in ErbB hyperactive tumor cells, the replication protein that is essential to viral replication will be released from plasma membrane and participate in virus replication, which leads to cell death and vector renewal.

ErbB-RASER delivered with adeno-associated virus (AAV) and bearing an apoptotic output (Bid) had been demonstrated to selectively eliminate ErbB-hyperactive cancer cells in culture. However, delivering RASER systems to all cancer cells would be infeasible with the non-replicating AAV, as that would require all cancer cells to have direct access to the compartment into which the virus is delivered (e.g. bloodstream or intraperitoneal space). This is implausible as even soluble antibodies cannot penetrate passively into tissues beyond 20 μm^27^. Absent transcellular transport mechanisms, calculations indicate that only < 50% of cells in a spherical tumor of diameter 200 μm would be accessible to AAV transduction, decreasing to < 10% of cells in a 1.2-mm mass. Even if all tumor cells had physical access to administered virus, infecting all tumor cells with AAV would require viral particle numbers in excess of normal cells in the shared delivery compartment. This would require immense amounts of virus and cause toxicity from the presence of viral proteins in normal cells. In contrast, mathematical modeling indicates that elimination of tumors by replicating viruses is possible^28^. Interestingly, modeling also suggests that elimination is more likely across different virus doses if additional cytotoxic treatments are co-administered^28^.

In this study, we report the engineering of therapeutic viral vectors that replicate within ErbB-hyperactive cancer cells with exquisite specificity, enabling local amplification of the vector within tumors to bypass the overdelivery requirement. Specifically, we use RASER to program the non-pathogenic vesicular stomatitis virus (VSV) to selectively replicate within and kill phosphoErbB-positive (pErbB(+)) cancer cells. The resulting VSV-ErbB-RASER contains only sequences of wild-type VSV plus RASER components to block replication in healthy cells, and so restricts rather than enhances VSV functionality, similar to approved replication-competent VSV-based vaccines^29,30^. VSV-ErbB-RASER decreased toxicity to normal cells by 10,000-fold compared to wild-type virus and more than 10-fold compared to VSV-ΔM51, an attenuated VSV similar to those currently in clinical trials. In mouse xenograft models of metastatic pancreatic and ovarian cancer, VSV-ErbB-RASER is both safer and more efficacious than VSV-ΔM51, confirming an improved therapeutic index *in vivo* and a mechanism of action independent of adaptive anti-tumor immunity. Finally, VSV-ErbB-RASER synergized with standard-of-care chemotherapy to dramatically reduce ovarian tumor growth and extend survival. Thus, an RNA virus engineered for specific tropism in ErbB-hyperactive cells may be a new type of therapeutic agent that can be effectively combined with standard-of-care chemotherapy to treat ErbB-driven metastatic carcinomas.

## Results

For the ErbB-RASER system to be an effective cancer therapy, a method to express the RASER proteins in most tumor cells *in vivo* is needed. To allow ErbB-RASER to be expressed in cancer cells not directly in contact with the delivery compartment, we hypothesized that we could use a replicative viral vector whose replication in turn is activated by the output of ErbB-RASER **(Fig. 1b)**. In this way, after an initially sparse infection into a pErbB(+) tumor, the RASER-expressing vector can propagate locally through the tumor. Ideally, this process would continue until most cancer cells are destroyed. In contrast, sparse transduction of normal tissue would not be followed by amplification, keeping toxicity low. This strategy is similar to earlier unrealized proposals for engineering cancer-specific viruses^31^, but contrasts with current virotherapy trials, which test viruses as pro-inflammatory agents to stimulate immune responses to tumor-associated antigens (TAAs)^32,33^.

### Engineering adenovirus for selective toxicity to pErbB(+) cells

For an initial test of this concept, we chose adenovirus-5 (Ad5) as a well characterized DNA virus. Ad5 replication proceeds through transcriptional cascades^34^, beginning with transcription of the Ad5 *E1A* gene by host RNA polymerase, followed by the *E1A* gene products activating further viral gene transcription^35^. The *E1A* promoter has previously replaced with cancer-inducible promoters to achieve preferential virus replication in cancer cells *in vitro*, but the resulting viruses were attenuated even in tumor cells^36,37^. The most clinically advanced Ad5 variants are instead deleted in the *E1B* gene. These variants preferential replication in some cancer cells^38^, but *E1B* deletion reduces toxicity to cancer cells compared to wild-type Ad5 as well^39^, and mechanisms for cancer preference remain incompletely understood and may vary across cancers^40^. Clinical experience with an *E1B*-deleted Ad5 approved in China for intratumoral injection^41^ suggests that immune activation contributes a major anti-tumor effect^42^.

We hypothesized that ErbB-RASER could robustly rewire hyperactive ErbB to Ad5 activation through control of *E1A* transcription via GAL4-UAS promoter sequences (**Supplementary Fig. 1a**). We constructed a membrane-tethered RASER substrate fusion with a synthetic GAL4-VP16 transcription factor as the effector at either the C-terminus or the N-terminus. Together with a RASER protease component, these substrate components created ErbB-RASER-C-GAL4-VP16 or ErbB-RASER-N-GAL4-VP16 systems, respectively **(Supplementary Fig. 1b)**. When these systems were expressed in human pancreatic cancer cells, GAL4-VP16 release occurred in BxPC3 cells, which are constitutively phosphoErbB-positive (pErbB(+)), but not in MIAPaCa2, which are phosphoErbB-normal (pErbB-normal). Of note, MIAPaCa2 cells express EGFR and HER2, but phosphorylation levels are low at baseline and are inducible by EGF stimulation, whereas BxPC3 cells constitutively exhibit high levels of phosphoHER2 that are not further EGF-inducible^43^. Thus, ErbB-RASER-N-GAL4-VP16 activation occurred in pancreatic cancer cells with constitutively active ErbB, but not in cells with normal ligand-dependent ErbB activity, similar to previous results with ErbB-RASER in breast cancer cells^25^. Furthermore, GAL4-VP16 release in BxPC3 cells did not occur if ErbB phosphorylation was inhibited by the ErbB-targeted tyrosine kinase inhibitor (TKI) lapatinib **(Supplementary Fig. 1b)**, indicating that effector release is pErbB-dependent. Thus, the ErbB-RASER system was readily adaptable to release GAL4-VP16 specifically in pErbB(+) cancer cells.

Next, we generated Ad5-ΔE3-ErbB-RASER-C virus, in which ErbB-RASER-C-GAL4-VP16 was expressed by a CMV promoter, the E1A promoter was replaced by GAL4-UAS, and the E3 gene was deleted for additional packaging capacity **(Supplementary Fig. 1c)**. Ad5-ΔE3-ErbB-RASER-N virus was constructed similarly with ErbB-RASER-N-GAL4-VP16 **(Supplementary Fig. 1c)**. We then characterized the pErbB-specificity and efficacy of these viruses, comparing with Ad5-ΔE3 parental virus in BxPC3 cells or MIAPaCa2 pancreatic cancer cells. As desired, Ad5-ΔE3-ErbB-RASER-N caused cytotoxicity in BxPC3 but not in MIAPaCa2 cells **(Supplementary Fig. 1d)**. Furthermore, adding lapatinib rescued BxPC3 cells from cytotoxicity, showing again that Ad5-ΔE3-ErbB-RASER-N replication was pErbB-dependent. Ad5-ΔE3 was toxic to both lines regardless of pErbB level, confirming that BxPC3 and MIAPaCa2 cells both support Ad5-ΔE3 replication. In contrast to Ad5-ΔE3-ErbB-RASER-N, Ad5-ΔE3-ErbB-RASER-C failed to ablate any cells. Potential reasons could be lower expression of the Ad5-ΔE3-ErbB-RASER-C components or the larger system size resulting in slower replication.

We next compared Ad5-ΔE3-ErbB-RASER-N to Ad5-ΔE3 in ability to suppress pErbB(+) cell growth. We applied viruses to BxPC3 or MIAPaCa2 pancreatic cancer cells, and to SKOV3 or MDAH2774 ovarian cancer cells. Ad5-ΔE3-ErbB-RASER-N suppressed growth of the pErbB(+) BxPC3 and SKOV3 cells, but had no noticeable effect on pErbB-normal MIAPaCa2 and MDAH2774 cells **(Supplementary Fig. 1e–g)**. Lapatinib restored SKOV3 and BxPC3 growth in the presence of Ad5-ΔE3-ErbB-RASER-N, again showing that growth suppression was dependent on ErbB phosphorylation, as desired. In contrast, Ad5-ΔE3 again reduced growth of all cell lines regardless of pErbB status, confirming these cells are susceptible to unregulated Ad5-ΔE3.

However, RASER-rewired Ad demonstrated suboptimal activity. Ad5-ΔE3-ErbB-RASER-N was less potent than Ad5-ΔE3 in killing and reducing growth of pErbB(+) BxPC3 and SKOV3 cells (**Supplementary Fig. 1d–f**), suggesting that its replication was slower than parental Ad5-ΔE3. This could be due to a delay in E1A transcription imposed by the need to transcribe RASER components, translate them, and then accumulate released GAL4-VP16 effector. Nevertheless, Ad5-ΔE3-ErbB-RASER-N validated the ability of ErbB-RASER to selectively sense ErbB hyperactivity when expressed from a replicating viral vector, motivating us to test vector types that can be more directly linked to RASER output.

### Engineering VSV for selective toxicity to pErbB(+) cancer cells

We hypothesized that a RASER-controlled vesicular stomatitis virus (VSV) may be able to ablate cancer cells more effectively than Ad5-ΔE3. VSV is a non-integrating RNA virus that features rapid replication^44^, large burst sizes^45^, and host cell killing by both apoptosis and necrosis^46,47^. Replication-competent VSV has been approved as a replication-competent vaccine vector^29,30^ and is currently in multiple clinical trials as cancer treatments ^48^. VSV encodes only five proteins in its 11-kb negative-strand RNA genome: nucleocapsid (N), phosphoprotein (P), matrix (M), glycoprotein (G) and the large (L) polymerase. The VSV G protein binds to low-density lipoprotein receptor and related receptors^49^, conferring broad tissue tropism in rodent and human tissues. However, to our knowledge VSV has not yet been engineered to respond to any synthetic regulation.

To rationally link ErbB hyperactivity to VSV replication, we developed an ErbB-RASER system that releases VSV P as the effector. P is essential for the activity of L and therefore indispensable for virus replication^50^. We first tested ErbB-RASER-P by lentiviral transduction, observing accumulation of released P in pErbB(+) BxPC3 cells but not pErbB-normal MIAPaCa2 cells **(Supplementary Fig. 2a,b)**. P accumulation was inhibited by lapatinib, indicating it required ErbB phosphorylation. To create the ErbB-rewired VSV, we then replaced the P gene in a YFP-expressing VSV genome (VSV-YFP, Indiana strain)^51^ with a dual expression cassette encoding the two ErbB-RASER-P proteins **(Supplementary Fig. 2c)**, creating VSV-ErbB-RASER1 virus.

We tested VSV-ErbB-RASER1 for the ability to suppress cell growth when inoculated sparsely with a multiplicity of infection (MOI) of 0.1. As desired, VSV-ErbB-RASER1 selectively replicated in pErbB(+) BxPC3 pancreatic cancer cells but not ErbB-normal MIAPaCa2 pancreatic cancer cells (**Supplementary Fig. 2d**), resulting in growth suppression specifically of the pErbB(+) BxPC3 cells (**Supplementary Fig. 2e**). Notably, VSV-ErbB-RASER1 is as effective as parental VSV-YFP or VSV-YFP-ΔM51 in inhibiting pErbB(+) BxPC3 growth (**Supplementary Fig. 2e**). Importantly, the ErbB kinase inhibitor lapatinib, which prevents ErbB autophosphorylation on cytosolic tyrosine residues, abolished VSV-ErbB-RASER1 replication and rescued cell numbers, confirming its pErbB-dependent mechanism of action, whereas lapatinib had no effect on VSV-YFP or VSV-YFP-ΔM51 (**Supplementary Fig. 2d.e**). Finally, VSV-ErbB-RASER1 was inactive in human MRC5 fibroblasts, while VSV-YFP replicated well and fully inhibited cell growth, and while VSV-YFP-ΔM51 showed intermediate levels of replication and cell growth suppression (**Supplementary Fig. 3a,b**).

We next investigated whether the pErbB-specificity of VSV-ErbB-RASER1 would generalize to breast and ovarian carcinomas, which frequently display constitutive ErbB2 (HER2) phosphorylation. In pErbB(+) SKOV3 ovarian cancer cells, VSV-YFP, VSV-YFP-ΔM51, and VSV-ErbB-RASER1 all replicated well and suppressed cell growth, but only VSV-ErbB-RASER1 was inhibited by lapatinib, as desired (**Supplementary Fig. 3a,b**). In addition, while VSV-YFP and VSV-YFP-ΔM51 inhibited growth of pErbB-normal MDAH2774 ovarian cancer cells, VSV-ErbB-RASER1 was completely inactive, also as desired (**Supplementary Fig. 3a,b**). VSV-ErbB-RASER1 also demonstrated pErbB-dependence in replication and cell growth suppression in pErbB(+) SKBR3 and pErbB-normal MCF7 breast cancer cells, although it was only able to suppress cell growth to approximately 50% of the extent of VSV-YFP and VSV-YFP-ΔM51 (**Supplementary Fig. 3a,b**).

Taken together, these results demonstrate that an ErbB-specific RASER system can efficiently prevent VSV replication in pErbB-normal cells while allowing VSV to suppress the growth of pErbB(+) cells.

### Safety and efficacy of VSV-ErbB-RASER1 *in vivo*

We hypothesized that VSV-ErbB-RASER1 can ablate tumor cells *in vivo* via direct killing of tumor cells without requiring the generation of adaptive immunity to tumor neoantigens. To prepare for *in vivo* testing, we first measured the maximum-tolerated dose (MTD) of VSV-YFP and VSV-YFP-ΔM51 in NOD *scid* gamma (NSG) mice **(Supplementary Fig. 4a)**. VSV-YFP delivered by intraperitoneal (i.p.) injection, a route under clinical testing for peritoneal metastatic cancer^32^, caused weight loss and lethality at a dose of 10^6^ TCID_50_ **(Supplementary Fig. 4a,b)**. Attenuated VSV-YFP-ΔM51 was tolerated at 10^9^ TCID_50_ but lethal at 10^10^ TCID_50_, while VSV-ErbB-RASER1 was tolerated at the higher dose of 10^10^ TCID_50_ **(Supplementary Fig. 4a,b)**. Based on these MTD results, VSV-ErbB-RASER1 is at least 10,000 times safer than VSV-YFP and 10 times safer than VSV-YFP-ΔM51.

Next, we evaluated the efficacy of VSV-ErbB-RASER1 against metastatic pancreatic cancer *in vivo.* As approximately 50% of pancreatic cancer patients present with i.p. metastases^48^, we used i.p. xenografts of pErbB(+) BxPC3 pancreatic cancer cells to model this situation **(Supplementary Fig. 5a)**. The use of NSG mice allows us to assess how well VSV can directly ablate tumor cells without anti-tumor adaptive immunity. One week after tumor implantation of luciferase-expressing BxPC3 cells, and every three weeks afterwards, mice received i.p. VSV-YFP-ΔM51 at its MTD (10^9^ TCID_50_), or VSV-ErbB-RASER1 at a matched vector dose, or VSV-ErbB-RASER1 at its MTD (10^10^ TCID_50_). The dosing interval was chosen to match that of the standard chemotherapy intervals with gemcitabine in humans, and follows previously published work in mice^52^.

Tumor imaging and survival analysis revealed that only VSV-ErbB-RASER1 improved outcomes in this metastatic pancreatic cancer model. Compared to mock treatment, VSV-ErbB-RASER1 at its MTD was associated with slower tumor growth **(Supplementary Fig. 5b)** and significantly longer survival **(Supplementary Fig. 5c)**. The observed median survival extension of 15 days or 30% (P = 0.0002) compares favorably to six days or 27% observed previously with gemcitabine^52^. In contrast, VSV-YFP-ΔM51 at its MTD was no better than mock treatment in tumor growth trajectories or survival (**Supplementary Fig. 5b,c**). Administering VSV-ErbB-RASER1 at the same dose as VSV-YFP-ΔM51 at MTD also did not significantly improve tumor growth or survival (P = 0.61).

Next, we compared VSV-ErbB-RASER1 to VSV-YFP-ΔM51 in a model of metastatic peritoneal ovarian cancer, which occurs in approximately 70% of ovarian cancer cases^53^. Human pErbB(+) SKOV3 ovarian cancer cells expressing luciferase were implanted i.p. in NSG mice, followed by the same treatment regimens as above **(Supplementary Fig. 6a)**. A three-week dosing interval was again chosen to match that of the standard of care, paclitaxel and carboplatin, in humans following a previously described mouse model^54^. We performed six cycles, as additional cycles of maintenance chemotherapy do not improve survival in patients, and treatment often stops before the sixth cycle if it is poorly tolerated or the cancer is non-responsive.

In this metastatic ovarian cancer model, we observed tumor growth was best suppressed by VSV-ErbB-RASER1 at its MTD, followed by lower-dose VSV-ErbB-RASER1 and then by VSV-YFP-ΔM51 at its MTD (**Supplementary Fig. 6b**). Treatment with either dose of VSV-ErbB-RASER1 significantly prolonged survival, with the higher dose extending median survival by 43 days or 150% compared to mock treatment (P = 0.0007, **Supplementary Fig. 6c**). In contrast, treatment with VSV-YFP-ΔM51 at its MTD did not produce a significant improvement in survival, only extending median survival by 8 days or 12% (P = 0.04). Together, these results show that restricting VSV replication to HER2-hyperactive cells can indeed produce a virotherapeutic that is safer *in vivo* than the widely tested ΔM51 variants of VSV, which then allows for higher achievable therapeutic efficacy.

### Rational improvement of VSV-ErbB-RASER potency

We sought to improve the potency of hyperactive ErbB-restricted VSV. In the above experiments, VSV-ErbB-RASER1 had similar per-dose activity as VSV-YFP-ΔM51 on pErbB(+) tumors *in vivo*, even if its higher MTD enabled higher achievable therapeutic efficacy. VSV-ErbB-RASER1 was also less efficacious than VSV-YFP-ΔM51 on pErbB(+) SKBR3 breast cancer cells *in vitro* **(Supplementary Fig. 3b)**. To improve RASER-gated VSV for better efficacy per dose unit, i.e. potency, we explored increasing expression of the RASER substrate component to release more P protein in pErbB(+) cancer cells. We encoded this component as a separate transcription unit upstream of the protease component, generating VSV-ErbB-RASER2 (**Fig. 2a**). To enhance noninvasive bioluminescence imaging of virus *in vivo*, we also replaced YFP with the orange fluorescent protein (OFP) mScarlet3 fused to NanoLuc. Finally, we replaced the YFP in VSV-YFP and VSV-YFP-ΔM51 with the OFP-NanoLuc fusion to create VSV-ON and VSV-ON-ΔM51 for matched comparisons (**Fig. 2a**).

**Fig. 2.**
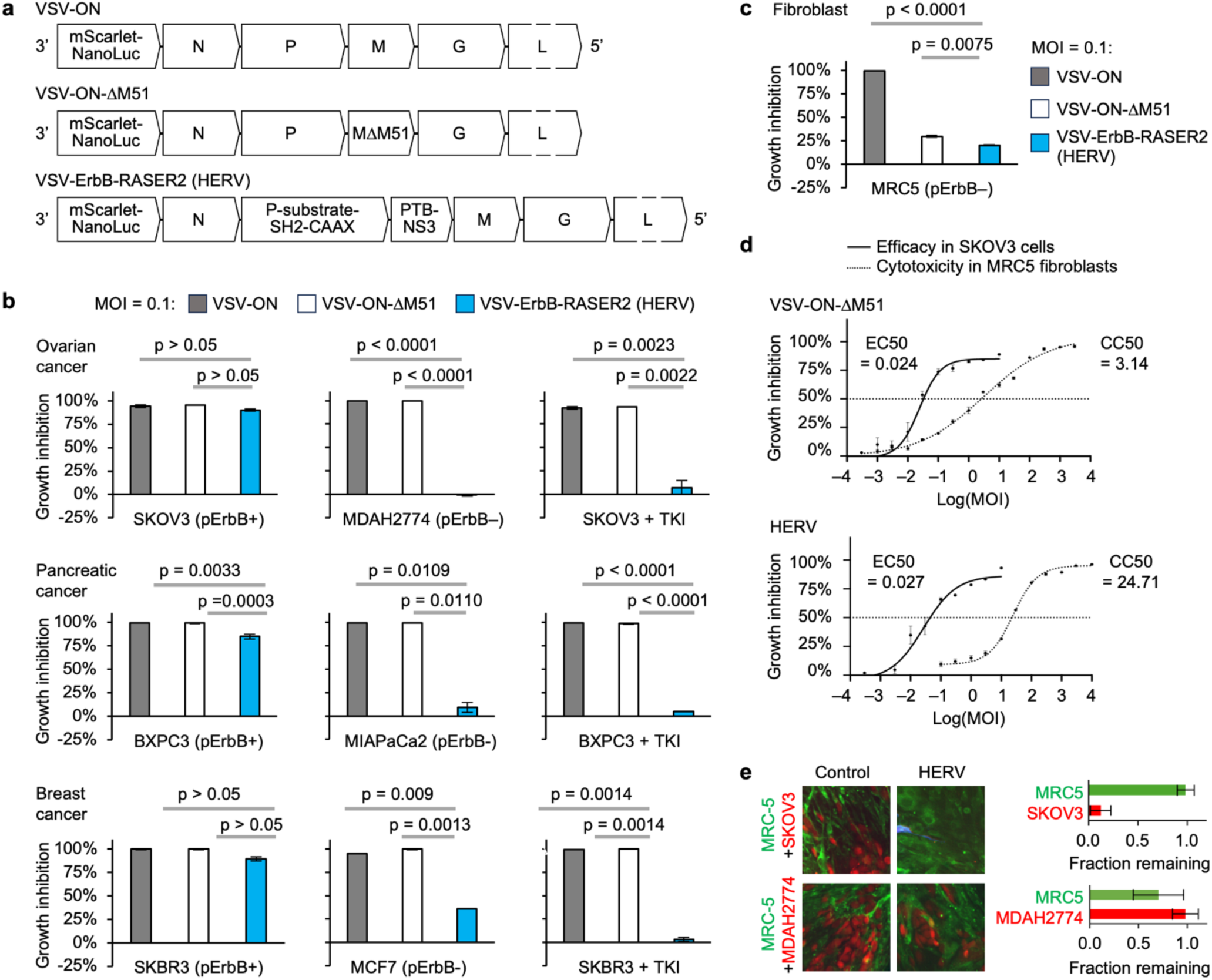
Specificity and potency of a hyperactive ErbB-restricted RNA vector (HERV). **a,** Genomic structure of VSV-ON, VSV-ON-ΔM51, and HERV. **b,** Growth inhibition by VSV-ON, VSV-ON-ΔM51, or HERV of ovarian cancer cells, pancreatic cancer cells, or breast cancer cells. Cells were infected with each vector at MOI of 0.1 and incubated with or without 0.5 μM of the TKI lapatinib. After 5 days, relative numbers of viable cells were measured by ATP-based bioluminescence assay. Error bars represent SEM from n = 3 infected wells. **c,** Growth inhibition by VSV-ON, VSV-ON-ΔM51, or HERV of MRC5 fibroblasts. Measurements were performed in parallel with those of **b** above. **d,** EC_50_ and CC_50_ measurements of HERV or VSV-ON-ΔM51. pErbB(+) SKOV3 cells and pErbB-normal MRC5 were infected with each vector at different MOIs. 4 days post-infection, relative numbers of viable cells were measured by ATP-based bioluminescence assay. EC_50_ on SKOV3 cells was 0.027 TCID50 for HERV and 0.024 TCID_50_ for VSV-ON-ΔM51. CC_50_ on MRC5 cells was 24.7 TCID_50_ for HERV and 3.14 TCID_50_ for VSV-ON-ΔM51. N = 9–12 biological replicates. **e,** HERV specificity in coculture of cancer cells and lung fibroblasts. MRC5 cells were cultured for two days, then overlaid with pErbB(+) SKOV3 or pErbB-normal MDAH2774 cells. One day later, cells were infected with HERV at MOI of 1. Error bars represent SEM from N = 3 infected wells.

We tested VSV-ErbB-RASER2 on multiple pErbB(+) carcinoma cells and type-matched pErbB-normal cells after sparse infection. VSV-ErbB-RASER2 suppressed growth of pErbB(+) BxPC3 pancreatic cancer, SKOV3 ovarian cancer, and SKBR3 breast cancer cells, but not of pErbB-normal type-matched MIAPaCa2, MDAH2774, and MCF7 cells **(Fig. 2b)**. Notably, VSV-ErbB-RASER2 was as effective as VSV-ON or VSV-ON-ΔM51 in inhibiting pErbB(+) SKBR3 growth, indicating that VSV-ErbB-RASER2 was indeed more potent than VSV-ErbB-RASER1. As desired, VSV-ErbB-RASER2 showed no inhibition of pErbB-normal MRC5 fibroblasts (**Fig. 2c**). Given the ability of VSV-ErbB-RASER2 to suppress a variety of pErbB(+) cancer cells with minimal toxicity to normal cells, we designated it as hyperactive ErbB-restricted virotherapeutic (HERV).

To rigorously quantify HERV specificity and potency, we measured CC_50_ (50% cytotoxic concentration on a normal cell) and EC_50_ (50% effective concentration on a cancer cell). These measurements are routinely performed for small-molecule anticancer drugs, but are lacking from most virotherapy studies. We measured CC_50_ in units of MOI for HERV and VSV-ON-ΔM51 in MRC5 lung fibroblasts and EC_50_ in SKOV3 ovarian cancer cells (**Fig. 2d**). CC_50_ was 7-fold higher for HERV than VSV-ON-ΔM51, demonstrating the improved specificity of HERV over VSV-ON-ΔM51. Most noticeably, partial growth suppression of normal cells at lower MOIs with VSV-ON-ΔM51 was greatly reduced with HERV. EC_50_ values on SKOV3 cells *in vitro* were similar between HERV and VSV-ON-ΔM51, resulting in a higher specificity index (CC_50_/EC_50_) for HERV (838 vs. 138). These results quantitatively demonstrate that the HERV has less toxicity to normal cells than VSV-ON-ΔM51.

We next asked if HERV could selectively remove pErbB(+) cells from a mixed culture of pErbB(+) and pErbB-normal cells. Indeed, HERV infection of a coculture of pErbB(+) SKOV3 ovarian cancer cells and MRC5 fibroblasts resulted in the selective loss of the SKOV3 cells **(Fig. 2e**). In contrast, HERV had no activity against type-matched pErbB-normal MDAH2744 cells cocultured with MRC5 cells.

Taken together, these results indicate that by rationally improving VSV-ErbB-RASER, we could achieve higher potency against pErbB(+) cancer cells while maintaining pErbB specificity, making the resulting HERV the preferred candidate for treating pErbB(+) tumors *in vivo*.

### Safety and efficacy of low-dose HERV monotherapy against metastatic ovarian cancer *in vivo*

To prepare for *in vivo* anti-tumor testing of HERV, we next compared its tropism and MTD with that of VSV-ON-ΔM51 in NSG mice with the same protocol as above **(Fig. 3a)**. Imaging of the red-shifted NanoLuc reporter revealed HERV to replicate to lower levels than VSV-ON-ΔM51 at each dose level (**Supplementary Fig. 7a**). As these mice do not bear tumor, this is consistent with the above *in vitro* finding that HERV is less active than VSV-ON-ΔM51 in normal cells. Most importantly, VSV-ON-ΔM51 caused rapid weight loss and death in most mice at doses of 3×10^9^ or 10^10^ TCID_50_, while HERV was tolerated at 3×10^9^ TCID_50_ and only lethal to a minority of mice at 10^10^ TCID_50_ **(Supplementary Fig. 7b, Fig. 3a)**. Thus, HERV is at least 3 times safer than VSV-ON-ΔM51 *in vivo*.

**Fig. 3.**
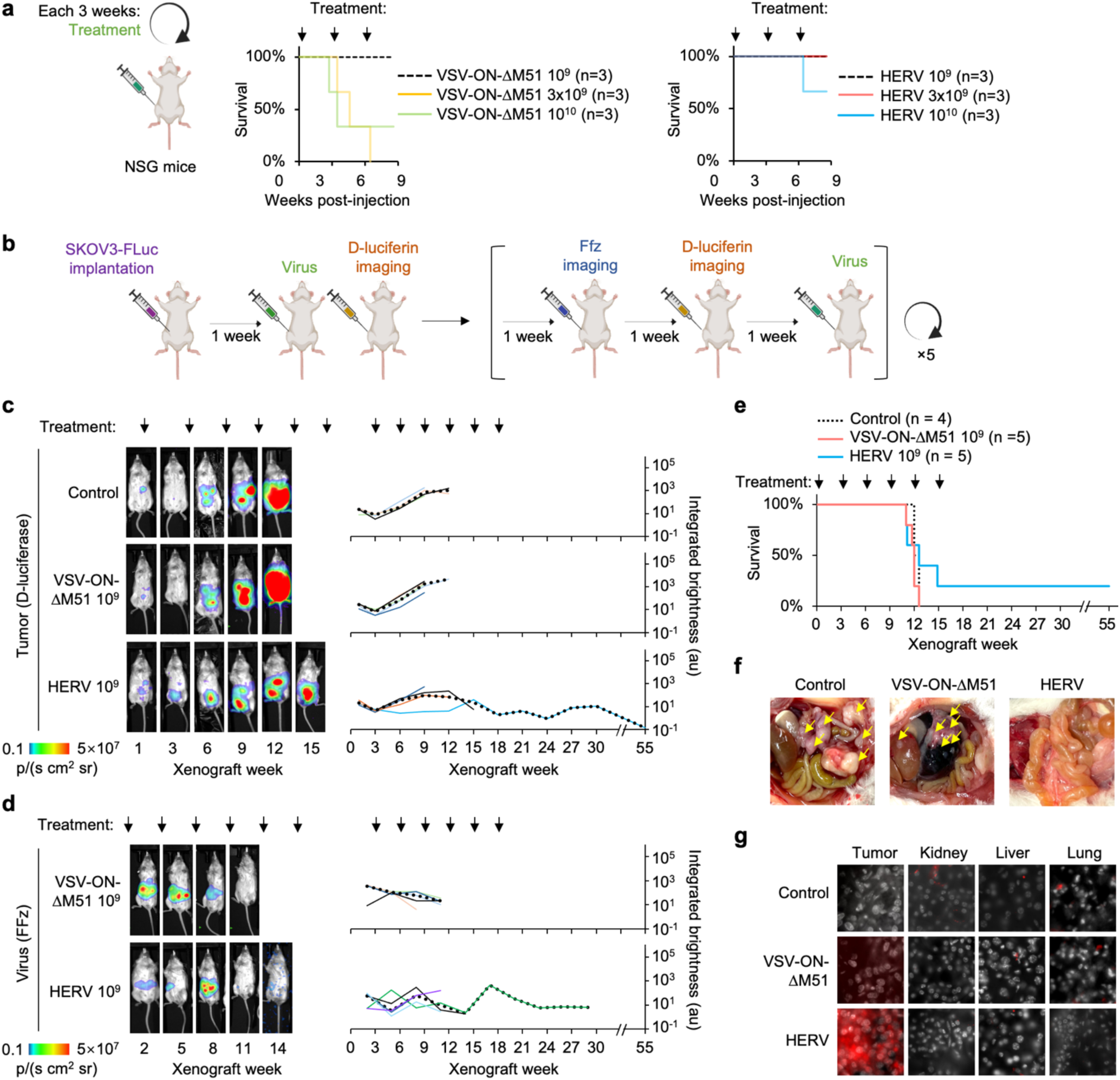
Efficacy of HERV monotherapy against metastatic ovarian cancer *in vivo*. **a,** Left, protocol to establish HERV MTD. 10^9^, 3×10^9^, or 10^10^ TCID_50_ of HERV or VSV-ON-ΔM51 were injected intraperitoneally (i.p.) into 2-month-old tumor-free female NSG mice every three weeks. Right, Kaplan-Meier (KM) survival curves. All mice were tested in parallel, but data are split into multiple charts for clarity, with the same control group shown in all charts. **b,** Mouse model of metastatic ovarian cancer. 10^6^ SKOV3 cells stably expressing FLuc were injected i.p. into 2-month-old NSG mice. Starting one week later, mice were treated by i.p. injection of PBS, 10^9^ TCID_50_ of VSV-ON-ΔM51, or 10^9^ TCID_50_ of HERV every 3 weeks. FFz and D-luciferin bioluminescence imaging was performed alternatively on the weeks the mice did not receive treatment. N = 4–5 mice per group. **c,** Bioluminescence imaging of SKOV3 tumor burden. **d,** Bioluminescence imaging of VSV-ON-ΔM51 and HERV reporter. In **c** and **d**, a mouse representative of the middle tertile from each group is shown and plotted as the thin black line. Dotted line represents mean bioluminescence from remaining mice. **e,** KM survival curves. Bonferroni-adjusted p values: 0.05 (mock vs. HERV 10^9^) by log-rank test. **f,** Pathology of mice assessed immediately after death. **g,** Viral reporter gene expression in tumor and organ sections.

We next compared the efficacy of HERV and VSV-ON-ΔM51 against metastatic peritoneal ovarian cancer *in vivo* **(Fig. 3b)**. NSG mice were implanted i.p. with pErbB(+) SKOV3 ovarian cancer cells expressing firefly luciferase (FLuc). One week after tumor implantation, and every three weeks afterwards for six treatment cycles, mice received i.p. VSV-ON-ΔM51 at its MTD (10^9^ TCID_50_) or HERV at the same dose, which is lower than its MTD. In each treatment cycle, tumor cells were imaged with D-luciferin and virus was imaged with fluorofurimazine (FFz). This *in vivo* efficacy comparison between HERV and VSV-ON-ΔM51 at the same dose (10^9^ TCID_50_) revealed HERV to be superior against ovarian cancer in multiple outcomes. First, HERV suppressed tumor growth compared to mock treatment, whereas VSV-ON-ΔM51 had no discernable effect **(Fig. 3c**). Second, HERV replicated better than VSV-ON-ΔM51 in the abdominal area of mice bearing i.p. SKOV3 cells, **(Fig. 3d**). Third, HERV improved survival duration over mock, whereas VSV-ON-ΔM51 had no discernable effect **(Fig. 3e)**. HERV also prevented the weight loss observed in mock-treated mice, whereas VSV-ON-ΔM51 again had no discernable effect (**Supplementary Fig. 8a**).

Consistent with the results of imaging, dissection of mice shortly after death revealed multiple large masses on peritoneal surfaces of control-and VSV-YFP-ΔM51-treated mice, whereas mice created with HERV showed no detectable nodules (**Fig. 3f**). Sectioning of organs and tumors showed that HERV was highly specific, replicating only in tumor tissue **(Fig. 3g)**.

In summary, HERV is at least 3 times safer than the attenuated VSV-ON-ΔM51and exhibits larger survival benefits in a mouse model of metastatic pErbB(+) ovarian cancer. Interestingly, one of the mice in the group receiving 10^9^ TCID_50_ HERV has remained tumor-free continuously for more than 9 months (**Fig. 3c**), surviving for more than 1 year after tumor implantation (**Fig. 3e**).

### HERV synergizes with chemotherapy to extend survival in metastatic ovarian cancer

We hypothesized that ErbB+ cancer cells in cell cycle arrest could still support HERV amplification and lysis, as VSV replication requires only host ribosomes and does no rely on deoxynucleotides or transcription factors whose abundance is modulated by the cell cycle. Previous studies showed that paclitaxel can stimulate cytoplasmic RNA virus replication, likely by inhibiting the expression of host antiviral genes^55^. Paclitaxel has also shown synergistic effects with Maraba virus in treating breast cancer both *in vitro* and *in vivo*^56^. Therefore, we hypothesized that HERV could synergize with the standard-of-care first-line therapy for metastatic ovarian cancer, paclitaxel and carboplatin doublet chemotherapy, to eliminate ovarian cancer cells *in vivo*.

We tested the efficacy of HERV in combination with paclitaxel and carboplatin on the same dosing schedule in the SKOV3 xenograft model of disseminated i.p. ovarian cancer. We compared HERV at its MTD, doublet chemotherapy, or the combination of HERV and doublet chemotherapy, administering therapy each three weeks and imaging bioluminescence from tumor and virus with each cycle (**Fig. 4a**). HERV or doublet chemotherapy each suppressed tumor growth relative to no treatment to a similar extent, with 0.71-log and 1.83-log reductions in median signal for VSV-ErbB-RASER and chemotherapy respectively at 9 weeks post-implantation, after which all the non-treated mice died (**Fig. 4b**). After 10 weeks, however, chemotherapy failed to prevent tumor growth.

**Fig. 4.**
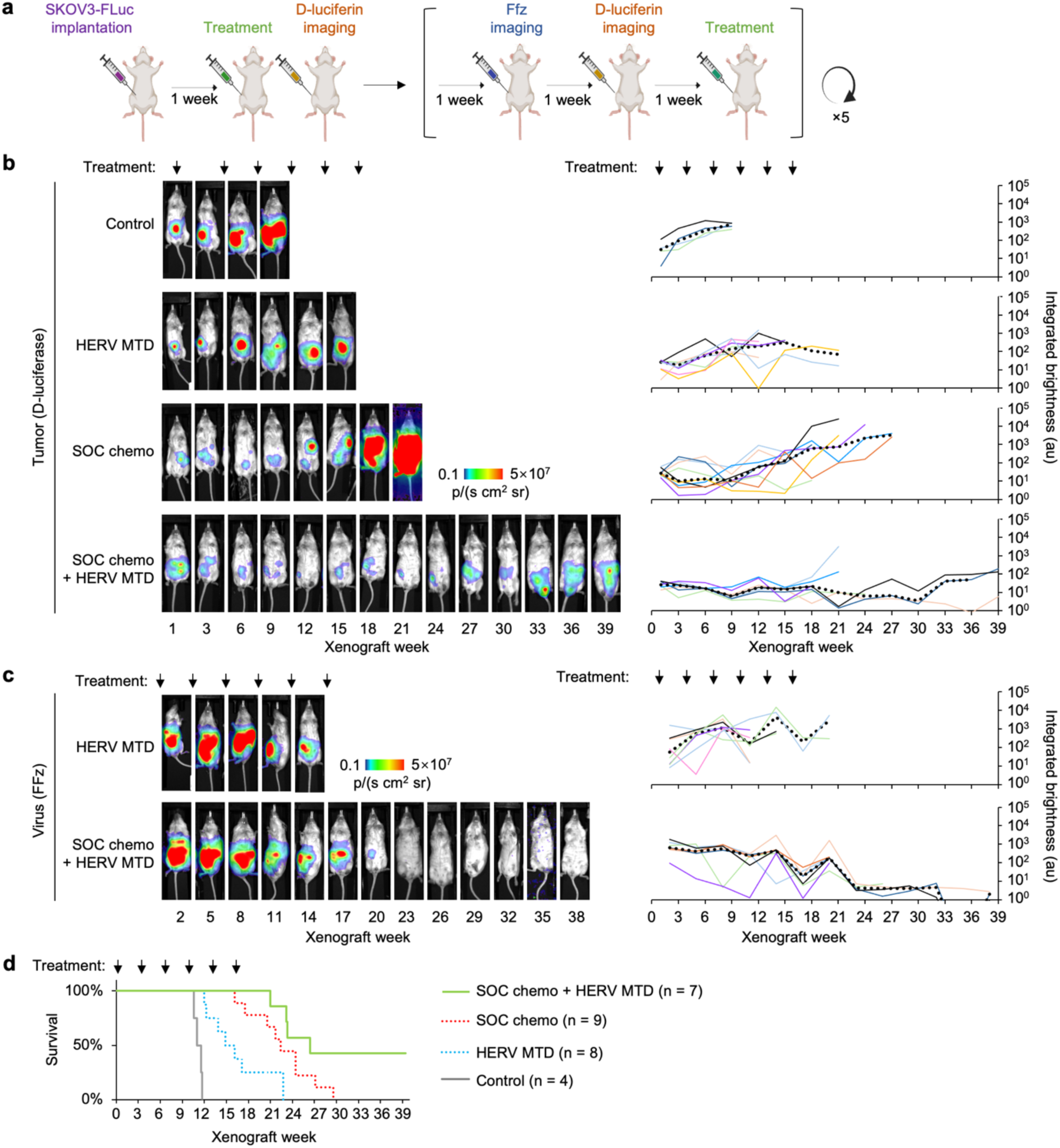
HERV synergizes with chemotherapy to extend survival in metastatic ovarian cancer. **a,** Mouse model of metastatic ovarian cancer. 10^6^ SKOV3 cells stably expressing FLuc were injected i.p. into 2-month-old NSG mice. Starting one week later, mice were treated by i.p. injection of vehicle, 3×10^9^ TCID_50_ (MTD) of HERV, standard-of-care (SOC) chemotherapy (40 mg/kg carboplatin and 10 mg/kg paclitaxel), or the combination of MTD HERV and SOC chemotherapy, every 3 weeks for 6 cycles. Bioluminescence imaging was performed with FFz or D-luciferin in the two non-treatment weeks. **b,** Bioluminescence imaging of SKOV3 tumor burden. **c,** Bioluminescence imaging of HERV virus replication. In **b** and **c**, a mouse representative of the middle tertile from each group is shown and plotted as the thin black line. Dotted line represents mean bioluminescence from surviving mice. **d,** Kaplan-Meier survival curves. Median survival in days: 77 for mock, 104 for MTD HERV, 157 for SOC chemotherapy, and 185 for the combination. Bonferroni-adjusted P values (log-rank test): <0.0001 mock vs. MTD HERV, <0.0001 mock vs. SOC chemotherapy, <0.0001 mock vs. combination, 0.0097 MTD HERV vs. SOC chemotherapy, 0.0001 MTD HERV vs. combination, and 0.0272 SOC chemotherapy vs. combination.

In contrast, the combination of HERV and chemotherapy caused levels of tumor-expressed FLuc to decrease continuously over time, including after 10 weeks (**Fig. 4b**). At 18 weeks, after treatment had ended, median tumor intensity in the combination treatment group was 0.73-log lower than in the remaining subjects of the HERV-only group, and 1.48-log lower than in the chemotherapy-only group, indicating strong synergy. In addition, after five treatment courses, two out of seven mice in the combination group showed undetectable levels of tumor cells, whereas tumor cells were continuously detectable in the individual treatment groups.

We also tracked HERV levels in the same experiment by imaging NanoLuc bioluminescence. Interestingly, viral levels demonstrated lower variability in the combination group compared to virus alone (**Fig. 4c**). This may be due to chemotherapy causing cell cycle arrest in tumor cells, preventing proliferation that can outrun viral replication. The absence of highly stochastic proliferative behavior may then result in more consistent levels of viral replication between subjects. Finally, the combination of HERV and chemotherapy did not cause more weight loss than HERV alone, indicating that toxicities were not additive (**Supplementary Fig. 8b**).

Most importantly, the combination of HERV with standard-of-care chemotherapy resulted in significantly greater median survival (185 days) than chemotherapy alone (157 days, p = 0.007) or HERV alone (104 days, p = 0.0001) (**Fig. 4d)**. Four months after the last treatment dose, all mice treated with chemotherapy alone or HERV alone had died, but 43% of mice treated with the combination were still surviving (**Fig. 4d**). In addition, the surviving mice maintained levels of tumor bioluminescence orders of magnitude lower than those observed in mice that had died (**Fig. 4b**). Taken together, these results indicate that HERV combines with doublet chemotherapy to synergistically ablate metastatic carcinoma *in vivo*, resulting in disease stabilization and dramatic prolongation of survival in a large fraction of subjects, which has not previously been observed in SKOV3 models of metastatic ovarian cancer.

## Discussion

In summary, we engineered viral vectors that express the EGFR or HER2-hyperactivity sensor ErbB-RASER, and whose replication in turn depends on ErbB-RASER output. ErbB-RASER effectively rewired oncogenic signaling to vector replication, resulting in selective ablation of ErbB(+) cancer cells. *In vivo*, VSV engineered to replicate only in response to the ErbB-RASER output significantly prolongs survival of NSG mice with metastatic pancreatic and ovarian disease, demonstrating that an oncogene-specific VSV can be effective as monotherapy without an anti-tumor immune response. Combination of ErbB specific VSV and standard-of-care chemotherapy successfully reduces tumor growth showing that synergy between VSV and chemotherapy could lead to better efficacy against metastatic carcinoma.

Our results demonstrate how rational engineering can widen the therapeutic window of a virus-based cancer therapy. Specifically, while VSV-YFP-ΔM51 did not extend survival in metastatic pancreatic and ovarian cancer models at its maximum tolerated dose, engineered HERV could be administered at a 3-fold higher dose, resulting in significant survival benefits. In addition, HERV could be combined with standard-of-care doublet chemotherapy without increasing any observed toxicity, while the combination exhibited clear synergy in ovarian tumor suppression *in vivo*. To our knowledge, HERV is the first example of programming a replicating therapeutic agent to specifically recognize pathological kinase signaling, as well as of restricting replication of a pure RNA virus to a defined biochemical signal.

HERV may also be the first case of a direct-acting tumoricidal agent that improves survival when added to carboplatin and paclitaxel doublet chemotherapy. The poly ADP-ribose polymerase (PARP) inhibitor olaparib has recently been approved for recurrent ovarian cancer, but adding it to doublet chemotherapy increases the frequency and severity of adverse events and interrupts chemotherapy dosing, even at only 8% of the standard dose^57^. Doublet chemotherapy is already dosed near maximum tolerated levels, with highly proliferative tissues such as the digestive and hematopoietic systems affected most severely, so it is not surprising that the addition of another proliferation-blocking agent further exacerbates toxicity^58^.

Currently the only agents with known benefits when added to doublet chemotherapy are the immune checkpoint inhibitor (ICI) antibodies such as anti-PD-L1 (programmed death-ligand 1)^59^ and bevacizumab^60^, whose immunity-enhancing and anti-angiogenesis mechanisms of action may generate toxicities that overlap less with chemotherapy. However, ICIs have their own severe adverse effects via the generation of autoimmune reactions. In addition, despite well established efficacy in EGFR-negative lung cancer^61^, ICIs produce response rates of <20% in most carcinomas, correlating with generally low tumor mutational burden (TMB)^62–64^. TMB measurements can identify a fraction of cases that are more likely to respond, but even this TMB-enriched response rate is still only 30%^65,66^. Thus, the majority of carcinoma patients, either low or high TMB, do not benefit from ICIs. In addition, even for lung cancer, median survival remains under 3 years after immunotherapy^61^.

Engineered replication-competent VSV vectors are currently being tested as cancer therapeutics, but current clinical candidates are also designed as immune-stimulating rather than tumor-ablating agents. For example, VSV-IFNβ-NIS (Voyager V1, VV1) is being tested as an immunostimulatory agent in combination with ICIs^67^. The production of IFNβ is intended to induce interferon-responsive genes in normal cells to protect against VSV-IFNβ-NIS replication^68^, while many cancer cells are defective in interferon signaling and would therefore permit VSV-IFNβ-NIS replication^69^. Subsequent lysis of the cancer cells by VSV-IFNβ-NIS would then cause inflammaion and promote neoantigen presentation, leading to adaptive immune responses against the tumor. Similarly the ΔM51 mutation in VSV-ΔM51 reduces the ability of M protein to suppress host cell NF-κB activation and IFNβ production^70^. However a large fraction of cancer cells, including ovarian cancer cells, are IFNβ-competent and resistant to VSV-ΔM51 or VSV-IFNβ-NIS, ironically requiring IFN-β pathway inhibition by ruxolitinb to achieve a therapeutic effect^71–73^. Regardless, the majority of carcinomas have low TMB and thus are poor targets for adaptive immunotherapy^62–64^. Thus, even if existing trials using VSV derivatives as stimulants for adaptive immunotherapy are successful, most carcinoma patients are unlikely to benefit, and will require new treatment options.

In this study, we selected experimental conditions to model clinical practice as closely as possible and allow comparisons to earlier studies. For the ovarian cancer model, we chose SKOV3 cells as they have been widely used in other studies. SKOV3 is believed to be an ovarian clear cell carcinoma (OCCC)^74^, which is more invasive than the more common high-grade serous carcinoma (HGSC)^75^, but both OCCC and HGSC frequently overexpress HER2 and cause disseminated disease^76–78^. We administered treatments with the typical chemotherapy interval of every 3 weeks, in contrast to earlier virotherapy studies that used more frequent dosing. Our choice of IP administration also reflects current evidence for its superior efficacy over intravenous administration for ovarian cancer chemotherapy^79^. In preclinical ovarian cancer studies, CAR-T cells are also more effective delivered i.p. than intravenously^80,81^, and IP CAR-T cells are currently in clinical trials (NCT05316129).

A limitation of using VSV in clinical applications is that neutralizing antibodies produced upon initial exposure could reduce the efficacy of repeated treatment^82^. However, unlike other non-integrating viruses such as measles, vaccinia, or adenovirus, only one viral protein, the glycoprotein, is exposed on the membrane surface. This should allow humoral immunity to be circumvented by altering the glycoprotein to serodistinct or poorly immunogenic homologues^83,84^. Drugs that suppress adaptive immunity such as cyclophosphamide have also been used to delay antibody-mediated viral clearance^85^. Here, the higher safety of VSV-ErbB-RASER in normal cells relative to earlier engineered VSV may provide a larger margin of safety, and immunosuppressive drugs can be withdrawn if adverse effects arise.

The development of resistance is a concern for any cancer therapy, but it is interesting to speculate whether the phenomenon of oncogene addiction may limit its occurrence with ErbB-RASER. Resistance to TKIs inevitably arises after a few months, typically from kinase domain mutations that reduce drug affinity or from activation of related RTKs^86^. Resistance to antibodies and CAR-T cells arises from antigenic escape, wherein mutations within the recognized antigen causes loss of binding^8,87–89^. Thus, pErbB(+) cancer cells typically develop resistance by restoring their original growth and survival signaling mechanisms rather than evolving new pathways, i.e. they are addicted to the original pathway^90^. However, pErbB(+) cancer cells cannot escape ErbB-RASER by upregulating ErbB signaling; rather that would render them more susceptible. To escape ErbB-RASER while continuing to grow and metastasize, they would have to evolve a delinking from ErbB-RASER while maintaining the activation of growth pathways by constitutive phosphoErbB. But as RASER uses the same pErbB-binding domains as the endogenous signaling effectors, this would require both mutating the ErbB oncogene to lose RASER connectivity and mutating signaling effectors to recognize the new ErbB variant, a difficult task.

## Outlook

Additional possibilities for increasing the safety and efficacy of VSV-RASER can be explored in the future. In case of toxicity, RASER output can be turned off with a clinically approved HCV protease inhibitor, as RASER output requires HCV protease activity. In addition, VSV-RASER remains compatible with strategies combining virotherapy with immunotherapy^91^. Indeed, the improved ability of VSV-RASER to ablate tumor cells compared to attenuated VSV would be expected to improve TAA recognition by any subsequent adaptive immune response. Notably, low therapeutic indices of previous viral vectors have made investigators reluctant to increase the strength of co-administered immunostimulants in regimens combining virotherapy and immunotherapy^32^. With its reduced propensity for off-tumor propagation, VSV-RASER could be expected to result in less autoantigen presentation compared to earlier VSV variants as well.

In conclusion, our results suggest a new molecular approach to treating cancer, in which synthetic proteins and engineered viruses identify and kill cells harboring specific oncogenic signals with high selectivity and autonomy. It will be interesting to determine which oncogenic signals can be detected and used to trigger cell ablation using this strategy. Finally, further studies are needed to determine how best to combine negative and positive immunomodulators with RASER-controlled VSV in the presence of a functional immune system.

## Methods

### DNA constructs

Plasmids encoding RASER cassettes were cloned by standard molecular biology techniques including polymerase chain reaction, restriction enzyme digestion, ligation, and homology-mediated assembly by In-Fusion enzyme (Clontech). All subcloned fragments were sequenced in their entirety to confirm successful construction. Full sequences of all plasmids used in this study are available upon request.

### Chemical reagents

Stock solutions of 500 μM lapatinib (TSZ Chem) in dimethylsulfoxide (Thermo Fisher) were stored at –20 °C. Thawed stock solutions were diluted into cell culture media to achieve a final treatment concentration of 500 nM. Carboplatin was freshly dissolved into Dulbecco phosphate buffered saline (DPBS, without calcium or magnesium, Corning 21−031-CV) to make 4mg/ml working solution. Paclitaxel was dissolved in 1:1 CremophorEL:ethanol to make a 6 mg/mL stock, which was diluted to 2 mg/mL in DPBS before injection.

### Antibodies

Primary antibodies and dilutions used for immunoblotting were: mouse monoclonal anti-V5 (Thermo, R960-25), 1:2000; rabbit monoclonal anti-β-actin (Abcam, ab8227), 1:2000; mouse anti-phosphoErbB2 (Tyr1248) (Cell Signaling Technology #2247). Secondary antibodies were LI-COR 680RD goat-anti-mouse, 680RD goat-anti-rabbit (LICOR Bio 926-68071), 800CW goat-anti-mouse (LICOR Bio 926-32210), and 800CW goat-anti-rabbit (LICOR Bio 926-32211), all used at 1:5000.

### Cell culture and transfection

BxPC3 (ATCC), MIAPaCa2 (ATCC) and SKOV3 (gift from H. Dai, Stanford University) cell lines were cultured at 37 °C in 5% CO_2_ in Roswell Park Memorial Institute 1640 medium (RPMI 1640, Thermo) supplemented with 10% fetal bovine serum (FBS, Thermo), and 100 U/mL penicillin and 100 μg/mL streptomycin (Life Technologies). HEK293T, BHK-VSV (BHK-21 cells stably expressing VSV-N, P and L proteins), MCF7 (gift from H. Chang, Stanford University), MRC5 (ATCC) and MDAH2774 (ATCC) cell lines were cultured at 37 °C in 5% CO_2_ in Dulbecco’s Modified Eagle’s Medium (DMEM, HyClone) supplemented with 10% FBS and 100 U/mL penicillin and 100 μg/mL streptomycin. SKBR3 (ATCC) cell line was cultured at 37 °C in 5% CO_2_ in McCoy’s 5A medium (ATCC) supplemented with 10% FBS and 100 U/mL penicillin and 100 μg/mL streptomycin. BHK-21 (ATCC) and BHK-T7 (RIKEN BRC) cell lines were cultured at 37 °C in 5% CO_2_ in Dulbecco’s Modified Eagle’s Medium (DMEM, HyClone) supplemented with 5% FBS and 100 U/mL penicillin and 100 μg/mL streptomycin. Cells were transfected using Lipofectamine 2000 (Thermo Fisher) in Opti-MEM (Thermo Fisher) according to the manufacturer’s recommended protocol.

### Virus genome construction, packaging and rescue, and purification

To rescue recombinant VSV, HEK293T cells were infected with vaccinia virus expressing T7 RNA polymerase (vTF-7, Imanis), and subsequently transfected with the full-length rVSV plasmid and plasmids with T7-driven VSV N, P, and L genes by Lipofectamine 2000 (Thermo). One day later, BHK-VSV cells were overlaid on top of the HEK293T cells. Three days post-transfection, supernatants were centrifuged and filtered through a 0.22-μm (pore size) filter and transferred onto fresh BHK-VSV cells. An alternative method was used for VSV-ErbB-RASER1-1 rescue. BHK-T7 cells were transfected with full-length rVSV plasmid and plasmids with T7-driven VSV N, P, and L genes using Lipofectamine 2000. 5–7 days post transfection, supernatants were collected, centrifuged and transferred onto fresh BHK-VSV cells for amplification. Recombinant VSV was concentrated and purified by Lenti-X concentrator (Takara) according to the manufacturer’s protocol. Titers of VSV stocks were determined by plaque assays on BHK-VSV cells. Recombinant adenoviruses were packaged by using the AdEasy system (Agilent). Briefly, 24 hours before transfection, 8×10^5^ HEK293A cells were seeded in 60mm plates. The next day, purified PacI-linearized recombinant Ad genome plasmid was transfected with Lipofectamine 3000 (Thermo). Cells were incubated for 2–3 weeks until massive cytopathic effects were observed, then collected and freeze-thawed 4 times at –80 °C and 37 °C. Then supernatants were collected, centrifuged and transferred onto fresh HEK293A cells for amplification.

### Immunoblotting

Cells in 24-well plates were washed twice in PBS, then lysed with 50 μL of 90 °C lysis buffer (100mM Tris HCl pH 8.0, 4% SDS, 20% glycerol, 0.2% bromophenol blue, 10% 2-mercaptoethanol), and sheared by sonication. After heating at 95°C for 2 minutes, cell lysates and Precision Plus dual-color standard (Bio-Rad) were loaded onto 4–12% NuPAGE bis-tris gels (Thermo Fisher). Gels were transferred to nitrocellulose membranes using a Trans-Blot Turbo Transfer System (Bio-Rad) and blocked with Superblock T20 blocking buffer (Thermo Fisher). Membranes were probed with primary antibodies followed by fluorophore-conjugated secondary antibodies, with washes after each step, all in Superblock T20 blocking buffer. Membranes were imaged using an Odyssey imaging system (LI-COR). Quantification of immunoblots was performed in ImageJ^92^.

### Cytotoxicity and viability assays

20,000 cells were seeded in 100 µL culture medium in a 96-well white microplate (Greiner Bio-One, Austria). 24 hours later, the culture medium was replaced with fresh culture medium with virus dilution with or without 0.5 μM lapatinib. For VSV infections, fluorescence was imaged after 3 days, and cell viability was determined after 5 days using the CellTiter Glo 2.0 kit (Promega, USA). For Ad infection, cell toxicity was determined using the CytoTox-Glo Cytotoxicity Assay Kit (Promega, USA) after 6 days, and cell viability was determined useding the CellTiter Glo 2.0 kit (Promega, USA) after 7 days. Bioluminescence was measured on a multi-mode microplate reader (Safire 2, TECAN).

### Coculture of ovarian cancer cells and lung fibroblasts

10^5^ MRC5 cells marked with CellTracker™ Deep Red Dye (Thermo) were seeded into glass-bottom 24-well plates. Two days later, 20,000 pErbB(+) SKOV3 or pErbB-normal MDAH2774 ovarian cancer cells marked with CellTracker™ Green CMFDA (Thermo) were overlaid. After 24 h of co-culture,HERV were infected into cells at MOI of 1. Viable cells were counted and expressed as a fraction of the number of cells in the negative uninfected control.

### Mice

Mouse procedures were performed in compliance with USDA and NIH ethical regulations and approved by the Stanford Institutional Animal Care and Use Committee. 2-month-old female NSG mice (Jackson Lab 005557) were used in all experiments. For the MTD assay, mice were injected intraperitoneally (i.p.) with 10^6^ or 10^7^ TCID_50_ of VSV-YFP, 10^9^ or 10^10^ TCID_50_ of VSV-YFP-ΔM51, 10^9^ or 10^10^ TCID_50_ of VSV-ErbB-RASER1, or PBS every 3 weeks. Weighing was performed 3 times weekly. Mice were monitored daily for weight and physical appearance, and euthanized at humane endpoints. For xenograft assays, BXPC3 stably transduced with Antares or SKOV3 stably transduced with firefly luciferase (FLuc) were used for xenograft implantation. 10^6^ cells were injected i.p. into 2-month-old female NSG mice (Jackson Lab, No. 005557). Starting 1 week later, mice were treated by i.p. injection of PBS, 10^9^ TCID_50_ of VSV-YFP-ΔM51, 10^9^ TCID_50_ of VSV-ErbB-RASER1, or 10^10^ TCID_50_ of VSV-YFP-ΔM51 every 3 weeks. Tumor growth was followed by weekly bioluminescence imaging by injecting the Nanoluc substrate fluorofurimazine (FFz) or FLuc substrate D-luciferin. Mice were mixed together before imaging, then selected at random for imaging. During imaging, the experimenter was blinded to treatment. Mice were anesthetized by inhalation of isoflurane for 5 min. For mice bearing BXPC3 xenografts expressing Nanoluc, 0.175 μmol of FFz and 0.8 mg poloxamer-407 (P-407) in 100 μL of DPBS was injected i.p. For mice bearing SK-OV-3 xenografts expressing FLuc, 3 mg D-luciferin were dissolved in 200 μL DPBS and injected i.p. Immediately after injection, bioluminescent images were acquired in an Ami HT system (Spectral Instruments Imaging) for 10 min at intervals of 1 min. Mouse index numbers were read from ear tags after each image was obtained. For figure preparation, cartoons were created with the assistance of BioRender.com.

### Quantification and statistical analysis

Statistical plans were formulated in consultation with the Stanford Biostatistics Shared Resource. For comparison of fibroblast growth suppression by VSV variants, a power analysis based on effect sizes in an exploratory experiment determined a two-fold difference could be detected with a sample size of 3 at a beta level of 0.8 and an alpha level of 0.05 in a single pairwise comparison by a two-tailed Student’s t test. The experiment was then performed with a single comparison of interest pre-determined for testing. Normality was assessed by the Shapiro-Wilk test. For survival analyses, a sample size of 8 mice was selected at the time of tumor implantation to match published statistical analyses of treatment effects on intraperitoneal ovarian xenografts. Pairwise Mantel-Cox log-rank tests were performed for all 6 comparisons between experimental groups, with the P value adjusted by the Bonferroni method for the number of comparisons. Alpha level for significance was 0.05.

## Acknowledgements

We thank Dr. H. K. Chung (University of North Carolina) for advice on RASER system adaptation, Dr. J. Sage (Stanford) for advice on tumor models, and Dr. Y. Su, Y. Wu, and L. Liu (Lin Lab, Stanford) for training on animal procedures and bioluminescence imaging.

## Funding

M.Z.L. discloses support for the research described in this study from the Alliance for Cancer Gene Therapy (Young Investigator Award) and the Damon Runyon Foundation (Damon Runyon-Rachleff Innovation Award). X.Z. discloses support from a Stanford Bio-X Fellowship and a Stanford Bioengineering Seibel Scholarship. K.T.B. discloses support from the National Institutes of Health (grant DP2AG067666). C.-Y.K. discloses support from the Canadian Institutes of Health Research (grant COV-440388).

## Author contributions

X.Z. designed and performed all experiments, analyzed data, prepared figures, and co-wrote the manuscript. K.T.B. and C.-Y.K. supplied previously unpublished reagents. M.Z.L. conceived general vector designs and experimental strategies, provided advice on experimental procedures and analysis, prepared figures, and co-wrote the manuscript.

## Competing interests

X.Z. and M.Z.L. are inventors on a patent application describing the viral constructs in this study.

---

**Supplementary Fig. 1.**
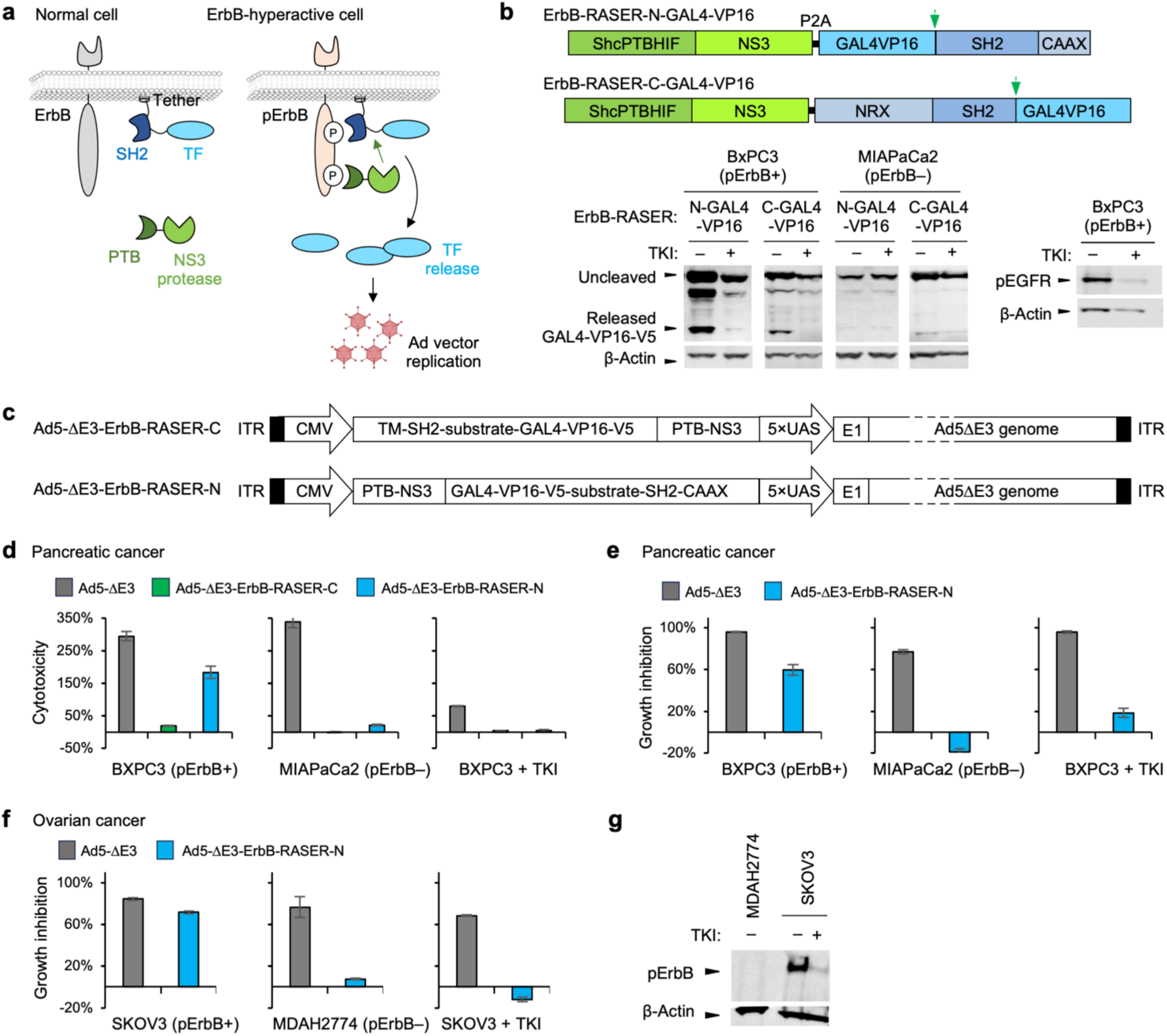
Engineering adenovirus for selective toxicity to pErbB(+) cancer cells. **a,** Strategy to link ErbB-RASER activation to Ad activation. TF, GAL4-VP16 transcription factor. **b,** GAL4-VP16 release efficiency from ErbB-RASER-N-GAL4-VP16 or ErbB-RASER-C-GAL4-VP16. BxPC3 and MIAPaCa2 cells were infected with lentivirus expressing each system at a multiplicity of infection (MOI) of 5. After 48 h with or without 0.5 μM lapatinib, cells were lysed for immunoblotting against the V5 epitope tag fused to GAL4-VP16. Lysates were also blotted with anti-phosphoEGFR antibody to validate the cell lines. β-actin served as a loading control. **c,** Genomic structure of Ad5-ΔE3-ErbB-RASER-C and Ad5-ΔE3-ErbB-RASER-N. The E1A promoter of Ad5-ΔE3 was replaced with a CMV-ErbB-RASER-C-GAL4-VP16 or CMV-ErbB-RASER-N-GAL4-VP16 transcription unit followed by a 5×UAS promoter. **d,** Cytotoxicity of Ad5 variants in pancreatic cancer cells with or without ErbB hyperactivity. BxPC3 and MIAPaCa2 cells were infected with Ad5-ΔE3-ErbB-RASER-C, Ad5-ΔE3-ErbB-RASER-N, or Ad5-ΔE3 at MOI of 10 and incubated with or without 0.5 μM of the ErbB TKI lapatinib. After 6 days, cytotoxicity was measured by CytoTox-Glo assay and plotted relative to noninfected control. Error bars represent standard error of the mean (SEM). n = 3 technical replicates. **e,f,** Potency comparison between Ad-ErbB-RASER-N and Ad5-ΔE3 in pancreatic and ovarian cancer cells. Cells were infected with Ad5-ΔE3-ErbB-RASER-N or Ad5-ΔE3 at the MOI of 10 and incubated with or without 0.5 μM lapatinib. After 7 days, viability was measured by ATP-based bioluminescence assay. Y axes represent growth inhibition, which equals 1 minus the relative viability to noninfected control. Error bars are SEM from n = 3 technical replicates. **g,** Anti-phosphoHER2 immunoblotting of MDAH2774 and SKOV3 cells was performed to confirm pErbB status. β-actin served as a loading control.

**Supplementary Fig. 2.**
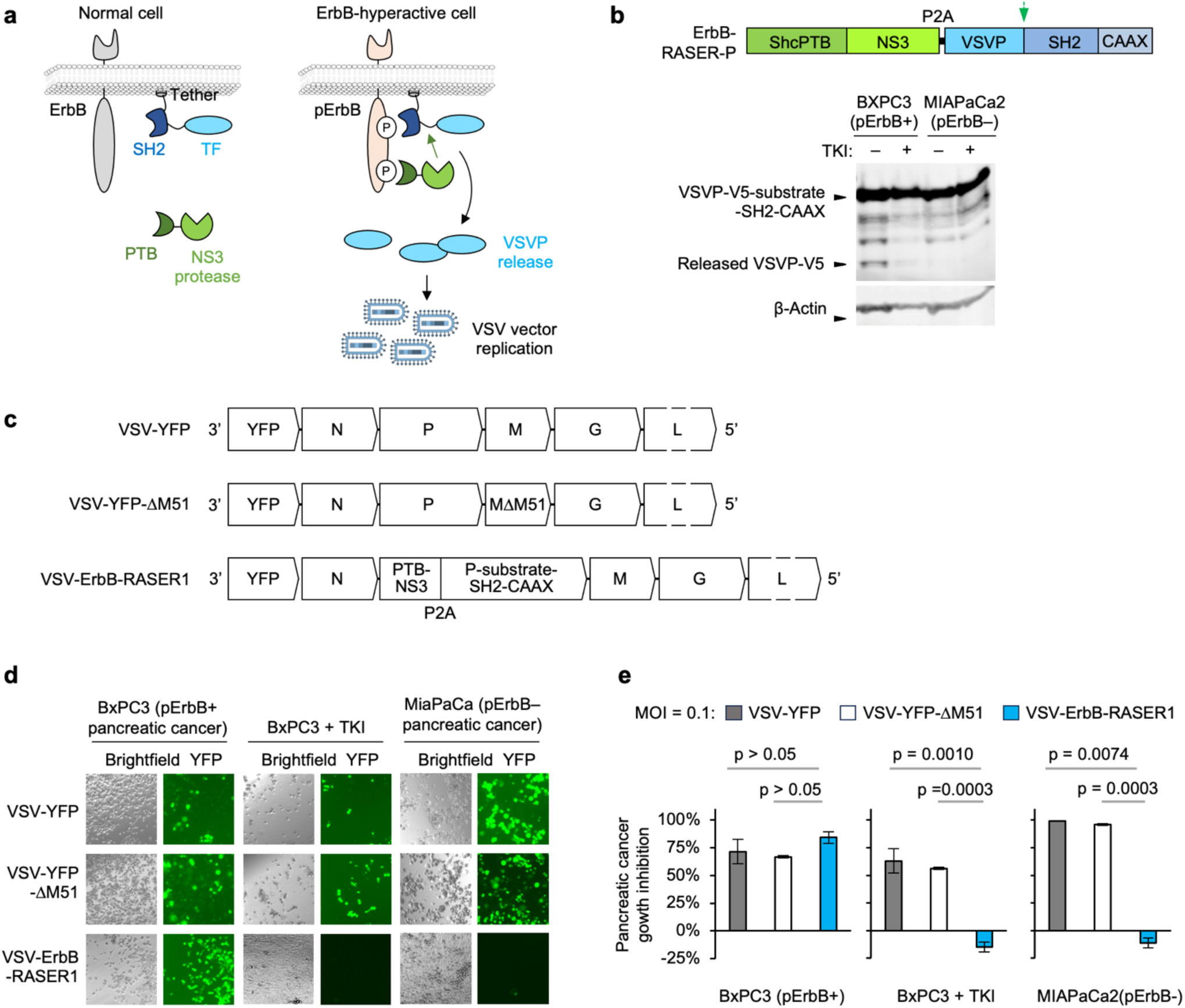
Engineering VSV for selective toxicity to pErbB(+) cancer cells. **a,** Strategy to link ErbB-RASER activation to VSV activation via release of VSV phosphoprotein (VSVP). **b,** Cleavage efficiency of VSV P from ErbB-RASER-VSVP (PTB-NS3 and VSVP-V5-substrate-SH2-CAAX) in pancreatic cancer cells by immunoblot. Experiments were performed as in Figure 2B. **c,** Genomic structure of VSV-YFP, VSV-YFP-ΔM51 and VSV-ErbB-RASER1. **d,** VSV-ErbB-RASER1 specificity and efficacy in pancreatic cancer cells. Cells were infected with VSV-YFP, VSV-YFP-ΔM51 and VSV-ErbB-RASER1 at MOI of 0.1 and treated with or without 0.5 μM of the ErbB kinase inhibitor lapatinib. Fluorescent images were taken 3 days post virus infection. VSV-ErbB-RASER1 replicated specifically in pErbB(+) BXPC3 cells, but not in pErbB-normald MIAPaCa2 cells. **e,** After 5 days, cell number reduction relative to uninfected controls were measured by ATP-based bioluminescence assay. Error bars represent SEM from n = 3 technical replicates. Scale bar represents 100 μm.

**Supplementary Fig. 3.**
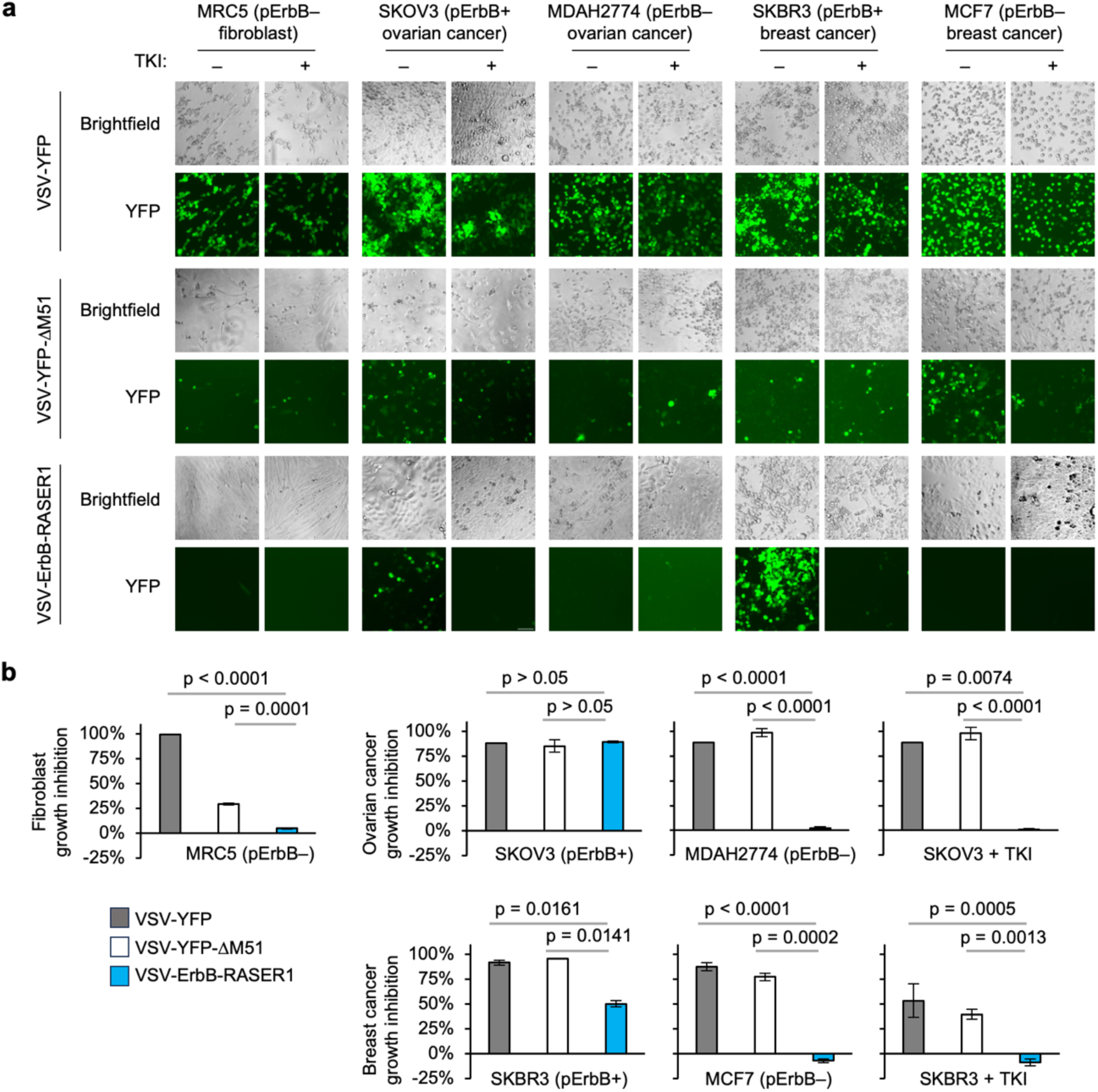
Specificity and efficacy of VSV-ErbB-RASER1 across tumor types. **a,** VSV-ErbB-RASER1 activity against in MRC5 fibroblasts, pErbB(+) and pErbB-normal ovarian cancer cells, and pErbB(+) and pErbB-normal breast cancer cells. Cells were infected with VSV-YFP, VSV-YFP-ΔM51, or VSV-ErbB-RASER1 at MOI of 0.1 and treated with or without 0.5 μM of the ErbB kinase inhibitor lapatinib. Images were taken 3 days post infection. Scale bar represents 100 μm. **b,** After 5 days, relative numbers of viable cells were measured by ATP-based bioluminescence assay. Error bars represent SEM from N = 3 infected wells. Scale bar represents 100 μm.

**Supplementary Fig. 4.**
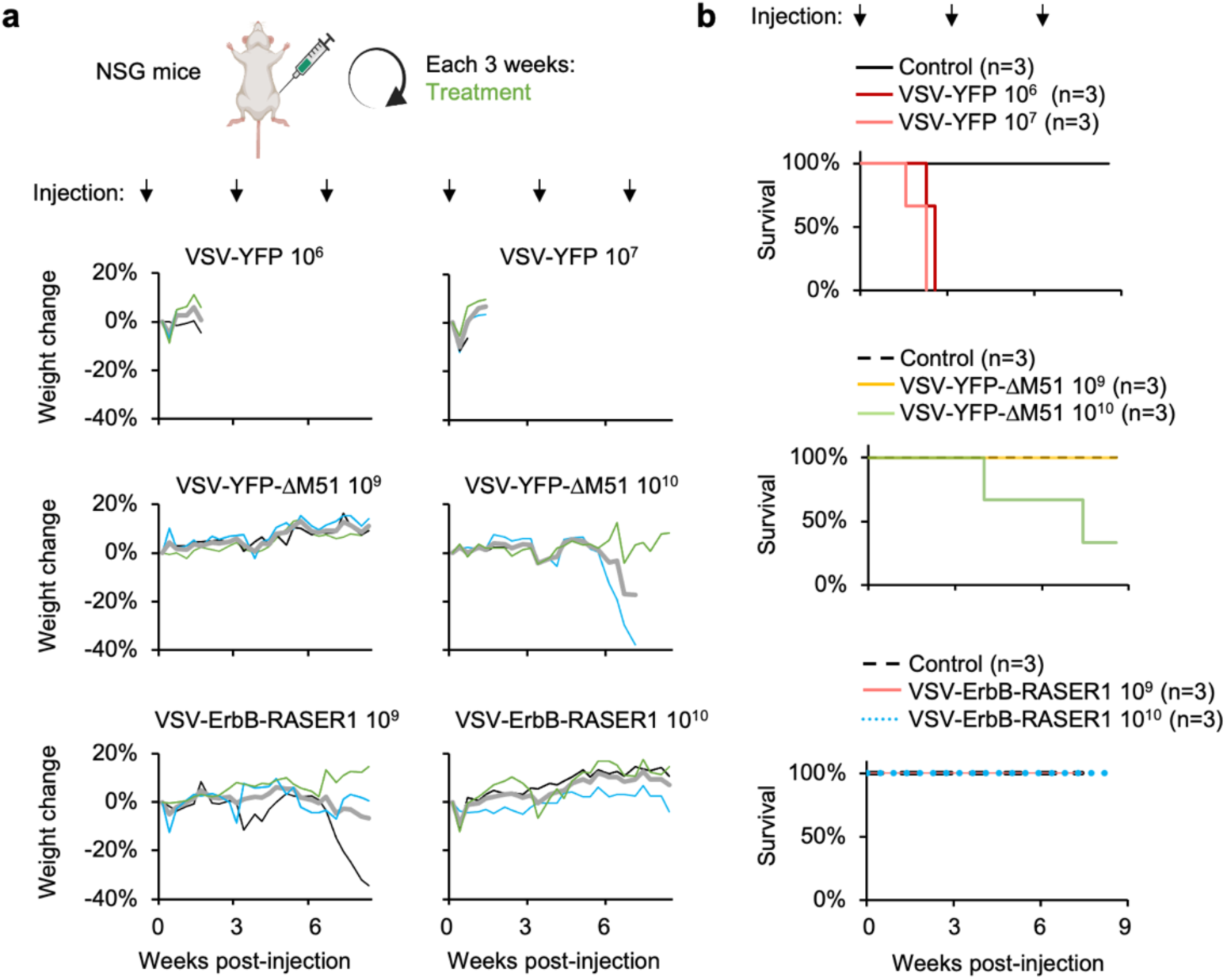
Safety of VSV-ErbB-RASER1 *in vivo*. **a,** Above, maximum-tolerated dose (MTD) study design. 10^6^ or 10^7^ TCID_50_ of VSV-YFP, 10^9^ or 10^10^ TCID_50_ of VSV-ErbB-RASER1, or PBS (control) were injected intraperitoneally (i.p.) into two-month-old female tumor-free NSG mice once every three weeks. Below, weights of mice over time in the six treatment groups. Lines represent individual mice. **b,** Kaplan-Meier survival curves for MTD determination. All mice were tested in parallel, but data are split into multiple charts for clarity, with the same control group shown in all charts.

**Supplementary Fig. 5.**
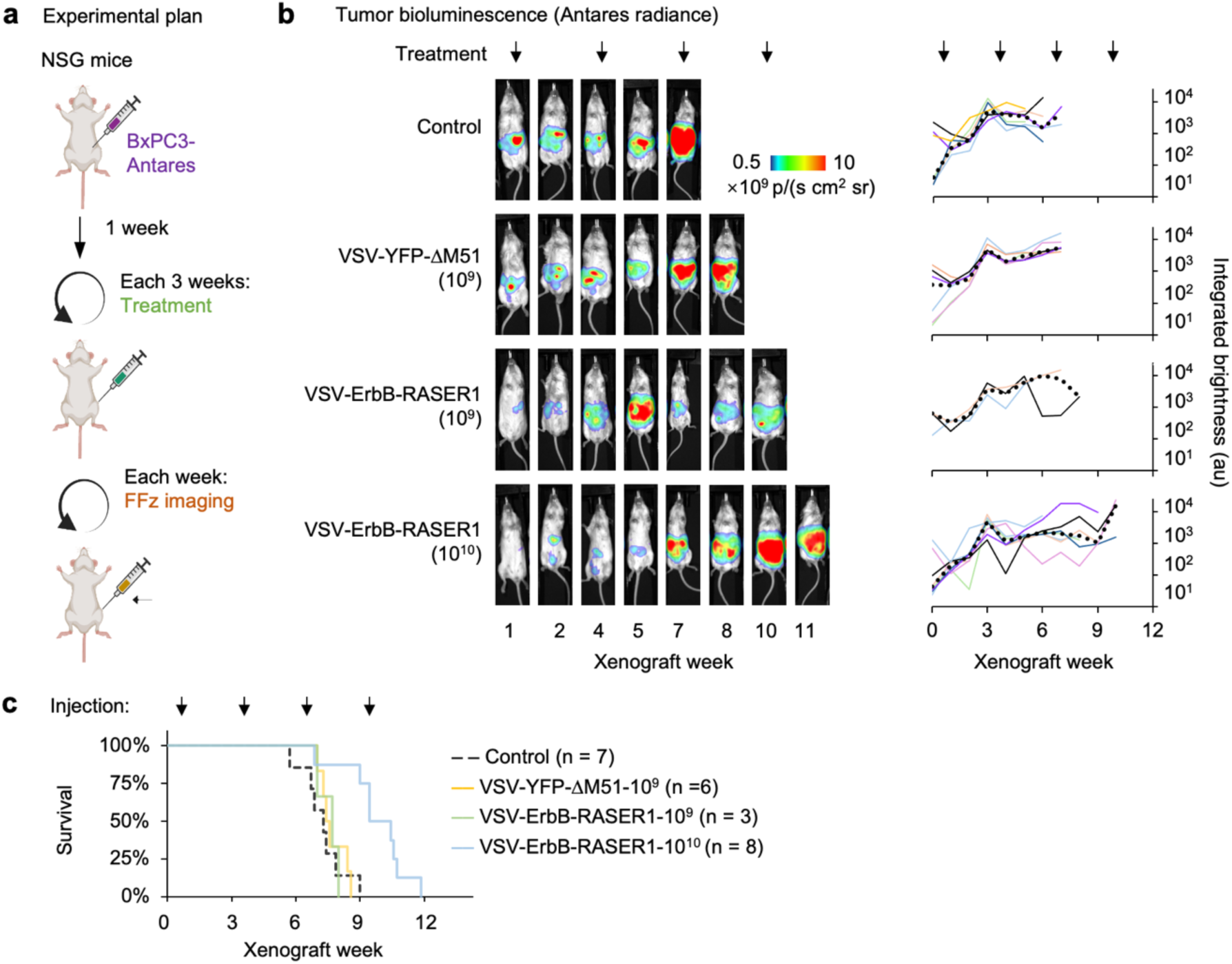
Efficacy of VSV-ErbB-RASER1 against metastatic pancreatic cancer *in vivo*. **a,** Mouse model of metastatic pancreatic cancer. 10^6^ BXPC3 cells stably expressing Antares luciferase were injected i.p. into 2-month-old NSG mice. Starting one week later, mice were treated by i.p. injection of PBS, 10^9^ TCID_50_ of VSV-YFP-ΔM51, 10^9^ TCID_50_ of VSV-ErbB-RASER1, or 10^10^ TCID_50_ of VSV-ErbB-RASER1 every 3 weeks. N = 3–8 mice per group. **b,** Bioluminescence imaging of BXPC3 tumor burden. Images of mice representative of the longest survivors from each group are shown, corresponding with thin black lines in the plots. Grey line represents average tumor bioluminescence readout from each group. **c,** Kaplan-Meier survival curves for efficacy determination. Median survival for mice was 51 days (mock), 52 days (VSV-YFP-ΔM51), 54 days (VSV-ErbB-RASER1 10^9^) and 66 days (VSV-ErbB-RASER1 10^10^). Bonferroni-adjusted p values: 0.61 (mock vs. VSV-YFP-ΔM51 10^9^), 0.61 (mock vs. VSV-ErbB-RASER1 10^9^), 0.0002 (mock vs. VSV-ErbB-RASER1 10^10^), 0.24 (VSV-YFP-ΔM51 10^9^ vs. VSV-ErbB-RASER1 10^9^), 0.0012 (VSV-YFP-ΔM51 10^9^ vs. VSV-ErbB-RASER1 10^10^), and 0.0025 (VSV-ErbB-RASER1 10^9^ vs. VSV-ErbB-RASER1 10^10^) by log-rank test.

**Supplementary Fig. 6.**
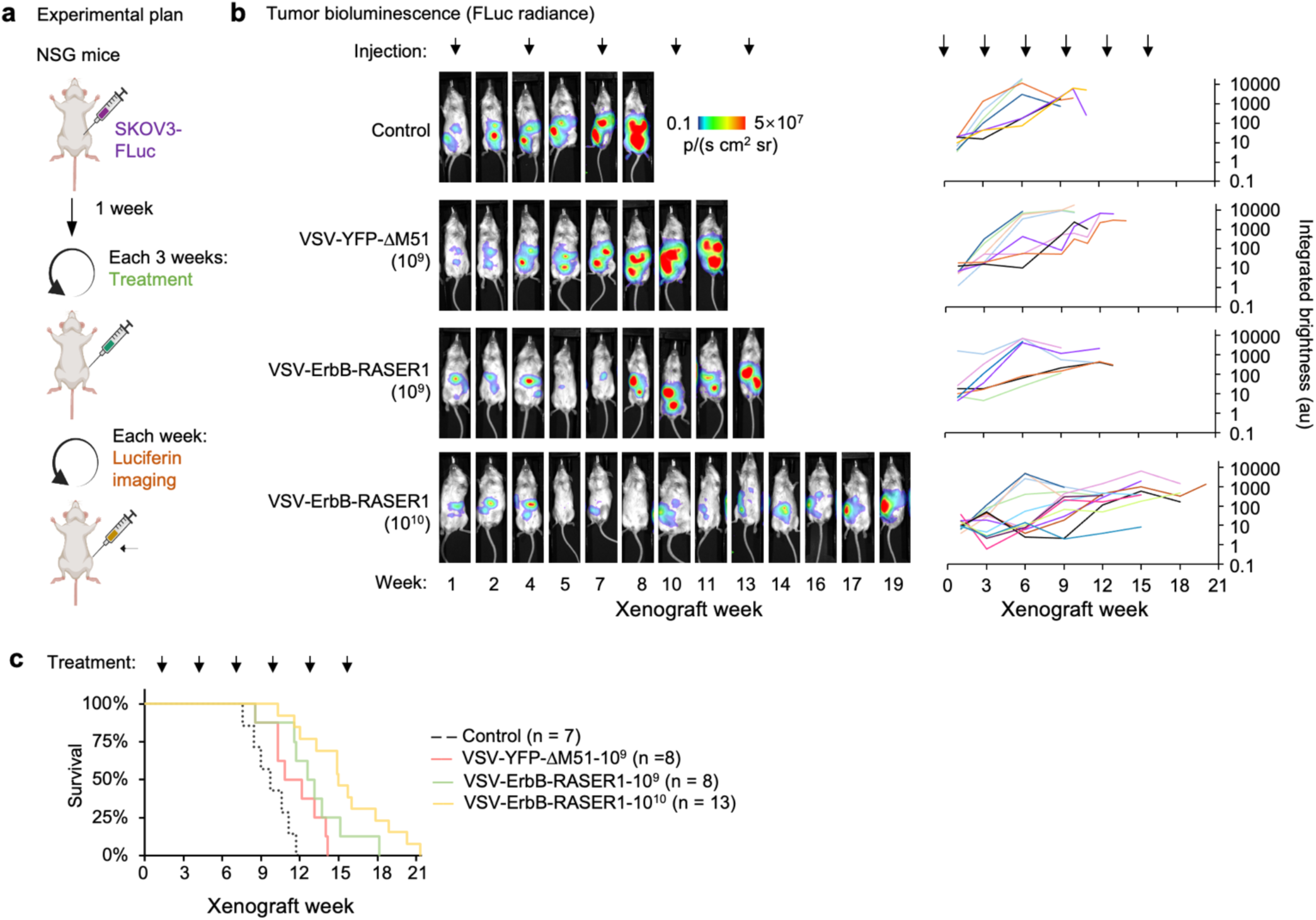
VSV-ErbB-RASER1 efficacy against metastatic ovarian cancer *in vivo*. **a,** Mouse model of metastatic ovarian cancer. 10^6^ SKOV3 cells stably expressing FLuc were injected i.p. into 2-month-old NSG female mice. Starting one week later, mice were treated by i.p. injection of PBS, 10^9^ TCID_50_ of VSV-YFP-ΔM51, 10^9^ TCID_50_ of VSV-ErbB-RASER1, or 10^10^ TCID_50_ of VSV-ErbB-RASER1 every 3 weeks. N = 7–13 mice per group. **b,** Bioluminescence imaging of SKOV3 tumor burden. A representative mouse from each group is shown. **c,** Kaplan-Meier survival curves for efficacy determination. Median survival was 68 days (mock), 76 days (VSV-YFP-ΔM51), 88 days (VSV-ErbB-RASER1 10^9^), and 105 days (VSV-ErbB-RASER1-10^10^). P values are 0.04 (mock vs. VSV-YFP-ΔM51 10^9^), 0.0009 (mock vs. VSV-ErbB-RASER1 10^9^), 0.0007 (mock vs. VSV-ErbB-RASER1-10^10^), 0.3 (VSV-YFP-ΔM51 10^9^ vs. VSV-ErbB-RASER1 10^9^), 0.009 (VSV-YFP-ΔM51 10^9^ vs. VSV-ErbB-RASER1 10^10^), and 0.08 (VSV-ErbB-RASER1 10^9^ vs. VSV-ErbB-RASER1 10^10^), compared to the Bonferroni-adjusted alpha level of 0.008.

**Supplementary Fig. 7.**
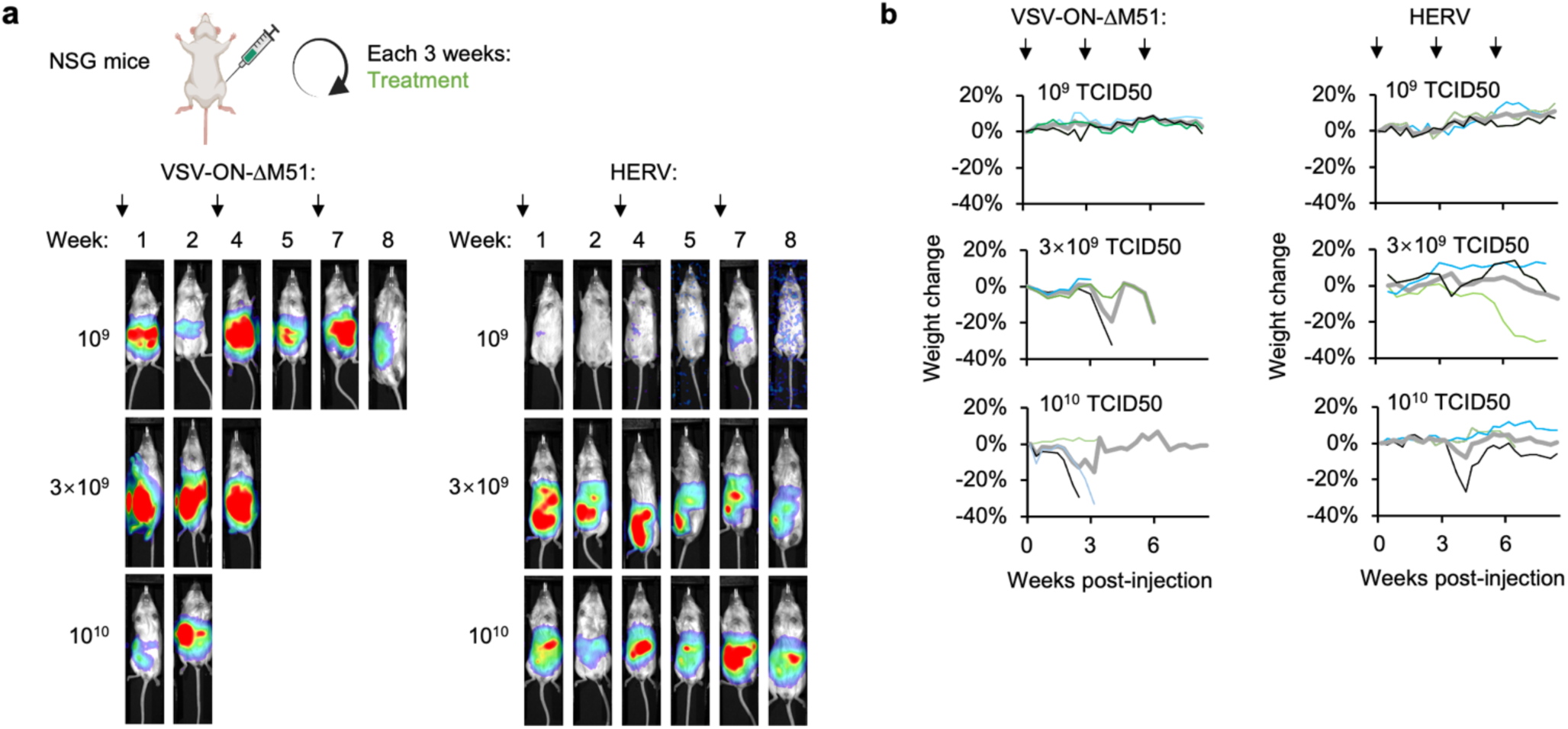
Safety of HERV *in vivo*. **a,** Bioluminescence readout of virus replication in the MTD assay performed with FFz. A representative mouse from each group is shown. **b,** Weight change of mice in the MTD assay. The black lines represent the mice shown in panel **a**. N=3 for each group.

**Supplementary Fig. 8.**
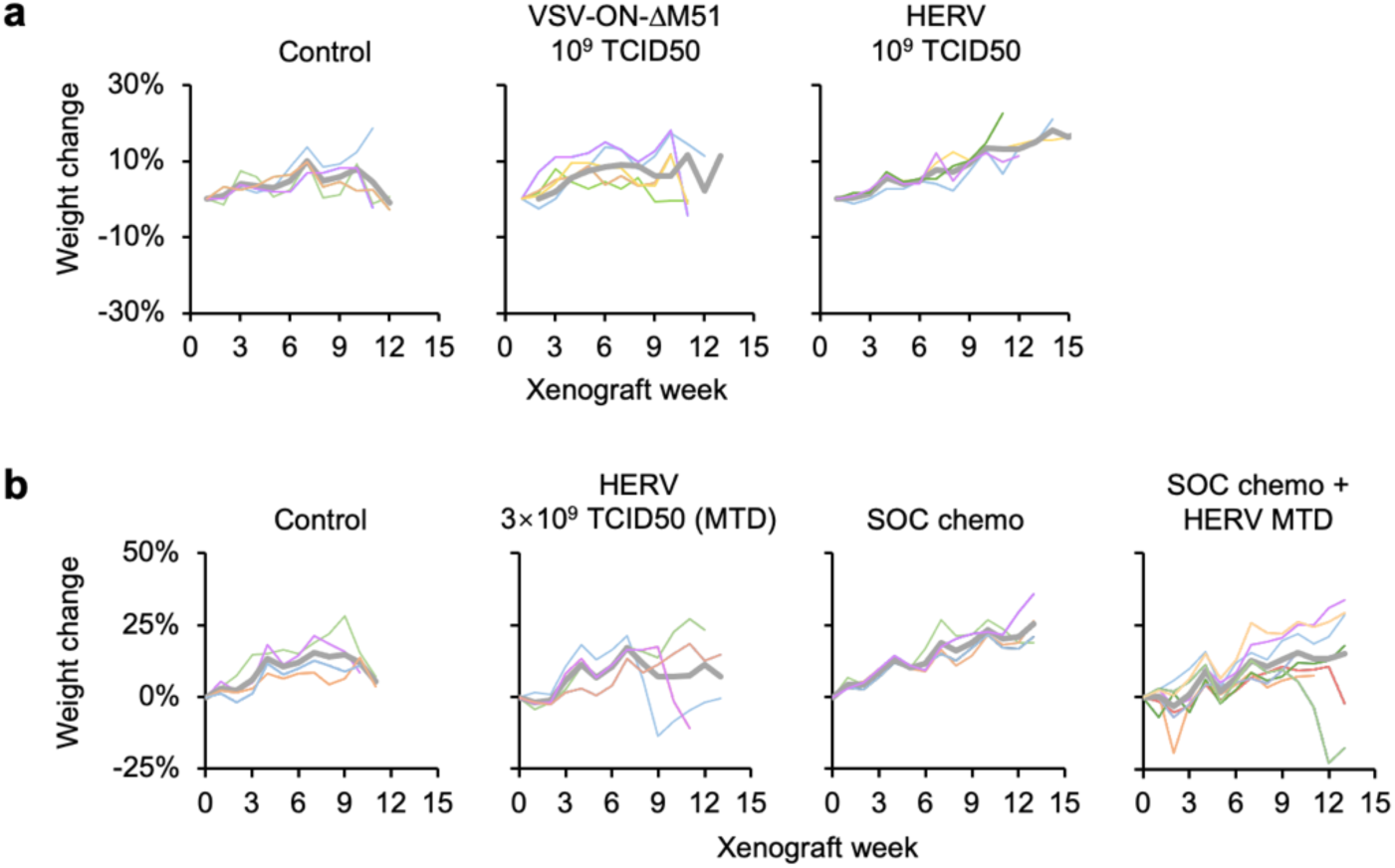
Weight changes during treatment for metastatic ovarian cancer. **a,** Weights of mice with SKOV3 ovarian cancer xenografts without treatment or with virotherapy at 10^9^ TCID_50_. Mice weight was monitored twice a week. 10^6^ SKOV3 cells stably expressing FLuc were injected i.p. into 2-month-old NSG female mice. Starting one week later, mice were treated by i.p. injection of PBS, 10^9^ TCID_50_ of VSV-ON-ΔM51, or 10^9^ TCID_50_ of HERV every 3 weeks. Mice received 6 doses of treatment in total. N = 4–5 mice per group. **b,** Weights of mice with SKOV3 ovarian cancer xenografts without treatment or with 3×10^9^ TCID_50_ of VSV-ErbB-RASER, chemotherapy, or combination therapy. N = 4–8 mice per group.

## Notes

### Summary of Updates

Figure 3 updated with recent results

## References

1. Bray, F., Laversanne, M., Weiderpass, E. & Soerjomataram, I. The ever-increasing importance of cancer as a leading cause of premature death worldwide. Cancer 127, 3029–3030 (2021).

2. Fares, J., Fares, M. Y., Khachfe, H. H., Salhab, H. A. & Fares, Y. Molecular principles of metastasis: a hallmark of cancer revisited. Signal Transduct Target Ther 5, 28 (2020).

3. Global Burden of Disease Cancer, C. et al. Cancer incidence, mortality, years of life lost, years lived with disability, and disability-adjusted life years for 29 cancer groups from 2010 to 2019: A Systematic Analysis for the Global Burden of Disease Study 2019. JAMA Oncol 8, 420–444 (2022).

4. Lin, J. J. et al. Five-year survival in EGFR-mutant metastatic lung adenocarcinoma treated with EGFR-TKIs. J Thorac Oncol 11, 556–565 (2016).

5. Jin, L. et al. Breast cancer lung metastasis: Molecular biology and therapeutic implications. Cancer Biol Ther 19, 858–868 (2018).

6. Torti, D. & Trusolino, L. Oncogene addiction as a foundational rationale for targeted anti-cancer therapy: promises and perils. Embo Molecular Medicine 3, 623–636 (2011).

7. Slamon, D. J. et al. Studies of the HER-2/neu proto-oncogene in human breast and ovarian cancer. Science 244, 707–712 (1989).

8. Mishra, R., Hanker, A. B. & Garrett, J. T. Genomic alterations of ERBB receptors in cancer: clinical implications. Oncotarget 8, 114371–114392 (2017).

9. Talia, K. L., Banet, N. & Buza, N. The role of HER2 as a therapeutic biomarker in gynaecological malignancy: potential for use beyond uterine serous carcinoma. Pathology 55, 8–18 (2023).

10. O’Shaughnessy, J., Gradishar, W., O’Regan, R. & Gadi, V. Risk of recurrence in patients with HER2+ early-stage breast cancer: Literature analysis of patient and disease characteristics. Clin Breast Cancer 23, 350– 362 (2023).

11. Cortés, J. et al. Trastuzumab deruxtecan versus trastuzumab emtansine in HER2-positive metastatic breast cancer: long-term survival analysis of the DESTINY-Breast03 trial. Nat Med (2024).

12. Schettini, F., Giudici, F. & Generali, D. Therapeutic resistance and optimal drug sequencing in HER2-positive metastatic breast cancer: unmet needs and future perspectives. Heliyon 10, e23367 (2024).

13. Swain, S. M., Shastry, M. & Hamilton, E. Targeting HER2-positive breast cancer: advances and future directions. Nat Rev Drug Discov 22, 101–126 (2023).

14. Scheck, M. K., Hofheinz, R. D. & Lorenzen, S. HER2-positive gastric cancer and antibody treatment: State of the art and future developments. Cancers (Basel) 16, 1336 (2024).

15. Meric-Bernstam, F. et al. Efficacy and safety of trastuzumab deruxtecan in patients with HER2-expressing solid tumors: Primary results from the DESTINY-PanTumor02 phase II trial. J Clin Oncol 42, 47–58 (2024).

16. Wen, M. et al. Combination of EGFR-TKIs with chemotherapy versus chemotherapy or EGFR-TKIs alone in advanced NSCLC patients with EGFR mutation. Biologics 12, 183–190 (2018).

17. Schlam, I. & Swain, S. M. HER2-positive breast cancer and tyrosine kinase inhibitors: the time is now. NPJ Breast Cancer 7, 56 (2021).

18. Skorda, A., Bay, M. L., Hautaniemi, S., Lahtinen, A. & Kallunki, T. Kinase Inhibitors in the Treatment of Ovarian Cancer: Current State and Future Promises. Cancers (Basel) 14, 6257 (2022).

19. Malaguti, P., D’Aloia, M. M. & Alimandi, M. The ErbB2 receptor in gastric cancer. the quick-change artist. Translational Gastrointestinal Cancer (2015).

20. Arienti, C., Pignatta, S. & Tesei, A. Epidermal growth factor receptor family and its role in gastric cancer. Frontiers in oncology (2019).

21. Doi, T. et al. Safety, pharmacokinetics, and antitumour activity of trastuzumab deruxtecan (DS-8201), a HER2-targeting antibody-drug conjugate, in patients with advanced breast and gastric or gastro-oesophageal tumours: a phase 1 dose-escalation study. Lancet Oncol 18, 1512–1522 (2017).

22. Shi, Y. et al. Efficacy, safety, and genetic analysis of furmonertinib (AST2818) in patients with EGFR T790M mutated non-small-cell lung cancer: a phase 2b, multicentre, single-arm, open-label study. Lancet Respir Med 9, 829–839 (2021).

23. Papini, F. et al. Hype or hope - Can combination therapies with third-generation EGFR-TKIs help overcome acquired resistance and improve outcomes in EGFR-mutant advanced/metastatic NSCLC? Crit Rev Oncol Hematol 166, 103454 (2021).

24. Wala, J. A. & Hanna, G. J. Chimeric antigen receptor T-cell therapy for solid tumors. Hematol Oncol Clin North Am (2023).

25. Chung, H. K. et al. A compact synthetic pathway rewires cancer signaling to therapeutic effector release. Science 364, eaat6982 (2019).

26. Waterman, H. & Yarden, Y. Molecular mechanisms underlying endocytosis and sorting of ErbB receptor tyrosine kinases. FEBS Lett. 490, 142–152 (2001).

27. Beltrán Hernández, I., et al. Imaging of tumor spheroids, dual-isotope SPECT, and autoradiographic analysis to assess the tumor uptake and distribution of different nanobodies. Mol Imaging Biol 21, 1079–1088 (2019).

28. Titze, M. I. et al. A generic viral dynamic model to systematically characterize the interaction between oncolytic virus kinetics and tumor growth. Eur J Pharm Sci 97, 38–46 (2017).

29. Garbutt, M. et al. Properties of replication-competent vesicular stomatitis virus vectors expressing glycoproteins of filoviruses and arenaviruses. J. Virol. 78, 5458–5465 (2004).

30. Anderson, E. M. & Coller, B.-A. G. Translational success of fundamental virology: a VSV-vectored Ebola vaccine. J. Virol. 98, e0162723 (2024).

31. Russell, S. J. & Peng, K. W. Measles virus for cancer therapy. Curr Top Microbiol Immunol 330, 213–241 (2009).

32. Harrington, K., Freeman, D. J., Kelly, B., Harper, J. & Soria, J. C. Optimizing oncolytic virotherapy in cancer treatment. Nat Rev Drug Discov 18, 689–706 (2019).

33. Lovatt, C. & Parker, A. L. Oncolytic viruses and immune checkpoint inhibitors: The “Hot” New Power Couple. Cancers (Basel) 15, (2023).

34. Nemerow, G. & Flint, J. Lessons learned from adenovirus (1970-2019). FEBS Lett. 593, 3395–3418 (2019).

35. Zheng, X. et al. Adenoviral E1a expression levels affect virus-selective replication in human cancer cells. Cancer Biol Ther 4, 1255–1262 (2005).

36. Huang, H. et al. Oncolytic adenovirus programmed by synthetic gene circuit for cancer immunotherapy. Nat Commun 10, 4801 (2019).

37. Liu, X. et al. Gene-viro-therapy targeting liver cancer by a dual-regulated oncolytic adenoviral vector harboring IL-24 and TRAIL. Cancer Gene Ther 19, 49–57 (2012).

38. Zhang, R., Cui, Y., Guan, X. & Jiang, X. A recombinant human adenovirus type 5 (H101) combined with chemotherapy for advanced gastric carcinoma: A retrospective cohort study. Front Oncol 11, 752504 (2021).

39. Bazan-Peregrino, M. et al. Comparison of molecular strategies for breast cancer virotherapy using oncolytic adenovirus. Hum Gene Ther 19, 873–886 (2008).

40. Cheng, P. H., Wechman, S. L., McMasters, K. M. & Zhou, H. S. Oncolytic replication of E1b-deleted adenoviruses. Viruses 7, 5767–5779 (2015).

41. Garber, K. China approves world’s first oncolytic virus therapy for cancer treatment. Journal of the National Cancer Institute 98, 298–300 (2006).

42. Zhang, Y. et al. Intraperitoneal oncolytic virotherapy for patients with malignant ascites: Characterization of clinical efficacy and antitumor immune response. Mol Ther Oncolytics 25, 31–42 (2022).

43. Larbouret, C. et al. In vivo therapeutic synergism of anti-epidermal growth factor receptor and anti-HER2 monoclonal antibodies against pancreatic carcinomas. Clin Cancer Res 13, 3356–3362 (2007).

44. Fields, B. N., Knipe, D. M. & Howley, P. M. Fields virology (Wolters Kluwer Health/Lippincott Williams & Wilkins, 2007).

45. Novella, I. S., Ball, L. A. & Wertz, G. W. Fitness analyses of vesicular stomatitis strains with rearranged genomes reveal replicative disadvantages. J. Virol. 78, 9837–9841 (2004).

46. Gaddy, D. F. & Lyles, D. S. Vesicular stomatitis viruses expressing wild-type or mutant M proteins activate apoptosis through distinct pathways. J. Virol. 79, 4170–4179 (2005).

47. Rogers, C. et al. Cleavage of DFNA5 by caspase-3 during apoptosis mediates progression to secondary necrotic/pyroptotic cell death. Nat Commun 8, 14128 (2017).

48. Lundstrom, K. Self-replicating vehicles based on negative strand RNA viruses. Cancer Gene Ther 30, 771– 784 (2023).

49. Finkelshtein, D., Werman, A., Novick, D., Barak, S. & Rubinstein, M. LDL receptor and its family members serve as the cellular receptors for vesicular stomatitis virus. Proc. Natl. Acad. Sci. U. S. A. 110, 7306–7311 (2013).

50. Das, S. C. & Pattnaik, A. K. Phosphorylation of vesicular stomatitis virus phosphoprotein P is indispensable for virus growth. J. Virol. 78, 6420–6430 (2004).

51. Beier, K. T. et al. Anterograde or retrograde transsynaptic labeling of CNS neurons with vesicular stomatitis virus vectors. Proc. Natl. Acad. Sci. U. S. A. 108, 15414–15419 (2011).

52. Maliandi, M. V. et al. AduPARE1A and gemcitabine combined treatment trigger synergistic antitumor effects in pancreatic cancer through NF-kappaB mediated uPAR activation. Mol Cancer 14, 146 (2015).

53. Mei, S., Chen, X., Wang, K. & Chen, Y. Tumor microenvironment in ovarian cancer peritoneal metastasis. Cancer Cell Int 23, 11 (2023).

54. Safra, T. et al. Weekly Carboplatin and Paclitaxel: A retrospective comparison with the three-weekly schedule in first-line treatment of ovarian cancer. Oncologist 26, 30–39 (2021).

55. Bressy, C., Droby, G. N. & Maldonado…, B. D. Cell cycle arrest in G2/M phase enhances replication of interferon-sensitive cytoplasmic RNA viruses via inhibition of antiviral gene expression. Journal of … (2019).

56. Bourgeois-Daigneault, M. C. et al. Combination of Paclitaxel and MG1 oncolytic virus as a successful strategy for breast cancer treatment. Breast Cancer Res 18, 83 (2016).

57. van der Noll, R. et al. Phase I study of intermittent olaparib capsule or tablet dosing in combination with carboplatin and paclitaxel (part 2). Invest. New Drugs 38, 1096–1107 (2020).

58. 58. Tang, H., et al. Estrone-conjugated PEGylated liposome Co-loaded paclitaxel and carboplatin improve anti-tumor efficacy in ovarian cancer and reduce acute toxicity of chemo-drugs. International Journal … (2022).

59. West, H., McCleod, M., Hussein, M. & Morabito…, A. Atezolizumab in combination with carboplatin plus nab-paclitaxel chemotherapy compared with chemotherapy alone as first-line treatment for metastatic non …. The Lancet … (2019).

60. Tewari, K. S., Burger, R. A. & Enserro…, D. Final overall survival of a randomized trial of bevacizumab for primary treatment of ovarian cancer. Journal of Clinical … (2019).

61. Ricciuti, B. et al. Association of high tumor mutation burden in non-small cell lung cancers with increased immune infiltration and improved clinical outcomes of PD-L1 blockade across PD-L1 expression levels. JAMA Oncol 8, 1160–1168 (2022).

62. Morand, S., Devanaboyina, M., Staats, H., Stanbery, L. & Nemunaitis, J. Ovarian Cancer Immunotherapy and Personalized Medicine. International Journal of Molecular Sciences 22, (2021).

63. Quintanilha, J. C. F. et al. Tumor mutational burden in real-world patients with pancreatic cancer: genomic alterations and predictive value for immune checkpoint inhibitor effectiveness. JCO Precis Oncol 7, e2300092 (2023).

64. Sha, D. et al. Tumor mutational burden as a predictive biomarker in solid tumors. Cancer Discov 10, 1808– 1825 (2020).

65. Kim, M. S. & Prasad, V. Pembrolizumab for all. J Cancer Res Clin Oncol 149, 1357–1360 (2023).

66. Marcus, L. et al. FDA Approval Summary: Pembrolizumab for the treatment of tumor mutational burden-high solid tumors. Clinical Cancer Research 27, 4685–4689 (2021).

67. Zhang, Y. & Nagalo, B. M. Immunovirotherapy based on recombinant vesicular stomatitis virus: Where are we? Front Immunol 13, 898631 (2022).

68. Jenks, N. et al. Safety studies on intrahepatic or intratumoral injection of oncolytic vesicular stomatitis virus expressing interferon-beta in rodents and nonhuman primates. Hum Gene Ther 21, 451–462 (2010).

69. Obuchi, M., Fernandez, M. & Barber, G. N. Development of recombinant vesicular stomatitis viruses that exploit defects in host defense to augment specific oncolytic activity. J. Virol. 77, 8843–8856 (2003).

70. Marquis, K. A., Becker, R. L., Weiss, A. N., Morris, M. C. & Ferran, M. C. The VSV matrix protein inhibits NF-κB and the interferon response independently in mouse L929 cells. Virology 548, 117–123 (2020).

71. Dold, C. et al. Application of interferon modulators to overcome partial resistance of human ovarian cancers to VSV-GP oncolytic viral therapy. Mol Ther Oncolytics 3, 16021 (2016).

72. Geoffroy, K., Mullins-Dansereau, V., Leclerc-Desaulniers, K., Viens, M. & Bourgeois-Daigneault, M.-C. Oncolytic vesicular stomatitis virus alone or in combination with JAK inhibitors is effective against ovarian cancer. Molecular Therapy: Oncology 32, 200826 (2024).

73. Patel, M. R. et al. JAK/STAT inhibition with ruxolitinib enhances oncolytic virotherapy in non-small cell lung cancer models. Cancer Gene Ther 26, 411–418 (2019).

74. Anglesio, M. S. et al. Type-specific cell line models for type-specific ovarian cancer research. PLoS One 8, e72162 (2013).

75. Hallas-Potts, A., Dawson, J. C. & Herrington, C. S. Ovarian cancer cell lines derived from non-serous carcinomas migrate and invade more aggressively than those derived from high-grade serous carcinomas. Sci Rep 9, 5515 (2019).

76. Koopman, T. et al. HER2 immunohistochemistry in endometrial and ovarian clear cell carcinoma: discordance between antibodies and with in-situ hybridisation. Histopathology 73, 852–863 (2018).

77. Woo, J. S. et al. Systematic assessment of HER2/neu in gynecologic neoplasms, an institutional experience. Diagn Pathol 11, 102 (2016).

78. Yeo, M. K., Kim, S., Yoo, H. J., Suh, K. S. & Kim, K. H. HER2 expression in peritoneal dissemination of high-grade serous ovarian carcinoma: A comparative study of immunohistochemical reactivity using four HER2 antibodies. J Clin Med 11, (2022).

79. Guchelaar, N. A. D. et al. Intraperitoneal chemotherapy for unresectable peritoneal surface malignancies. Drugs 83, 159–180 (2023).

80. Deshet-Unger, N. et al. Comparing intraperitoneal and intravenous personalized ErbB2CAR-T for the treatment of epithelial ovarian cancer. Biomedicines 10, 2216 (2022).

81. Domínguez-Prieto, V. et al. Understanding CAR T cell therapy and its role in ovarian cancer and peritoneal carcinomatosis from ovarian cancer. Front Oncol 13, 1104547 (2023).

82. Roberts, A., Buonocore, L., Price, R., Forman, J. & Rose, J. K. Attenuated vesicular stomatitis viruses as vaccine vectors. J. Virol. 73, 3723–3732 (1999).

83. Bishnoi, S., Tiwari, R., Gupta, S., Byrareddy, S. N. & Nayak, D. Oncotargeting by vesicular stomatitis virus (VSV): Advances in cancer therapy. Viruses 10, (2018).

84. Muik, A. et al. Pseudotyping vesicular stomatitis virus with lymphocytic choriomeningitis virus glycoproteins enhances infectivity for glioma cells and minimizes neurotropism. J. Virol. 85, 5679–5684 (2011).

85. Qiao, J. et al. Cyclophosphamide facilitates antitumor efficacy against subcutaneous tumors following intravenous delivery of reovirus. Clinical Cancer Research 14, 259–269 (2008).

86. Marrocco, I. & Yarden, Y. Resistance of lung cancer to EGFR-specific kinase inhibitors: Activation of bypass pathways and endogenous mutators. Cancers (Basel) 15, 5009 (2023).

87. Reslan, L., Dalle, S. & Dumontet, C. Understanding and circumventing resistance to anticancer monoclonal antibodies. MAbs 1, 222–229 (2009).

88. Jin, S. et al. Emerging new therapeutic antibody derivatives for cancer treatment. Signal transduction and targeted therapy 7, 39 (2022).

89. Mishra, A., Maiti, R. & Mohan…, P. Antigen loss following CAR-T cell therapy: Mechanisms, implications, and potential solutions. European Journal of … (2024).

90. Petroni, G., Buque, A., Coussens, L. M. & Galluzzi, L. Targeting oncogene and non-oncogene addiction to inflame the tumour microenvironment. Nat Rev Drug Discov 21, 440–462 (2022).

91. Shalhout, S. Z., Miller, D. M., Emerick, K. S. & Kaufman, H. L. Therapy with oncolytic viruses: progress and challenges. Nat Rev Clin Oncol 20, 160–177 (2023).

92. Schneider, C. A., Rasband, W. S. & Eliceiri, K. W. NIH Image to ImageJ: 25 years of image analysis. Nat Methods 9, 671–675 (2012).

